# Modulation of GABA_A_ receptor trafficking by WWC2 reveals class-specific mechanisms of synapse regulation by WWC family proteins

**DOI:** 10.1101/2024.03.11.584487

**Authors:** Thomas L. Dunham, Julia R. Wilkerson, Richard C. Johnson, Richard L. Huganir, Lenora J. Volk

## Abstract

WWC2 (WW and C2 domain-containing protein) is implicated in several neurological disorders, however its function in the brain has yet to be determined. Here, we demonstrate that WWC2 interacts with inhibitory but not excitatory postsynaptic scaffolds, consistent with prior proteomic identification of WWC2 as a putative component of the inhibitory postsynaptic density. Using mice lacking WWC2 expression in excitatory forebrain neurons, we show that WWC2 suppresses GABA_A_R incorporation into the plasma membrane and regulates HAP1 and GRIP1, which form a complex promoting GABA_A_R recycling to the membrane. Inhibitory synaptic transmission is dysregulated in CA1 pyramidal cells lacking WWC2. Furthermore, unlike the WWC2 homolog KIBRA (WWC1), a key regulator of AMPA receptor trafficking at excitatory synapses, deletion of WWC2 does not affect synaptic AMPAR expression. In contrast, loss of KIBRA does not affect GABA_A_R membrane expression. These data reveal unique, synapse class-selective functions for WWC proteins as regulators of ionotropic neurotransmitter receptors and provide insight into mechanisms regulating GABA_A_R membrane expression.

## INTRODUCTION

The WW and C2 domain-containing (WWC) family of proteins (WWC1-3) are associated with human cognition and implicated in pathologies of the central nervous system (CNS). Single-nucleotide polymorphisms (SNPs) in *WWC1* (aka KIBRA, KIdney/BRAin protein) are associated with variation in normal human memory performance ^1–8^ as well as risk for schizophrenia (SZ), Tourette (TS), and Alzheimer’s disease (AD) ^9–12^. Increased *WWC2* gene expression is associated with SZ risk and decreased *WWC2* gene expression is reported in Huntingtin’s disease patients ^13,14^. Additionally, copy number variations in *WWC2* have been reported in the MSSNG database of *de novo* and heritable cases of autism spectrum disorder ^15^. Reduced *WWC3* expression has been observed in glioma tissue and has been identified as a candidate marker of major depressive disorder (MDD) ^16,17^.

Despite the relevance of all WWC family proteins to disordered brain function, KIBRA is the only WWC family member that has been the subject of mechanistic study in the brain. KIBRA functions as an excitatory synapse scaffold, regulating activity-dependent trafficking of α-amino-3-hydroxy-5-methyl-4-isoxazolepropionic acid receptors (AMPARs), which mediate the majority of fast excitatory synaptic transmission in the brain ^18–21^. Deletion of KIBRA causes aberrant trafficking and expression of AMPAR subunits GluA1 and GluA2 in the hippocampus and impairs synaptic plasticity and memory ^18,19,21^. Additionally, loss of KIBRA impairs experience-induced changes in hippocampal and cortical circuit dynamics that support memory consolidation ^22^

WWC2 shares approximately 50% residue identity with KIBRA (human WWC2 vs. KIBRA: 49.37%, mouse WWC2 vs. KIBRA: 48.87%) and contains protein-protein and protein-lipid interacting domains common to the WWC family: two N-terminal WW domains, a C2 domain, an atypical protein kinase C (aPKC) binding site, and a C-terminal PDZ ligand ^23,24^ (Fig. 1A). In spite of its high degree of homology with KIBRA, which is localized at excitatory postsynaptic sites ^19^, WWC2 was identified as a putative member of the inhibitory postsynaptic density (iPSD) via proximity ligation proteomics ^25^. Specifically, WWC2 was labeled by each of the three BiRA-conjugated inhibitory postsynapse proteins investigated (Gephryin, InSyn1, and Arhgef9), and showed high fold enrichment on par with known iPSD proteins ^25^. Based on the homology of WWC2 to KIBRA and its putative interactions with inhibitory synaptic proteins, we hypothesized that WWC2 may play a role in trafficking neurotransmitter receptors at inhibitory synapses, complementing the role that KIBRA plays at excitatory synapses.

**Figure 1.**
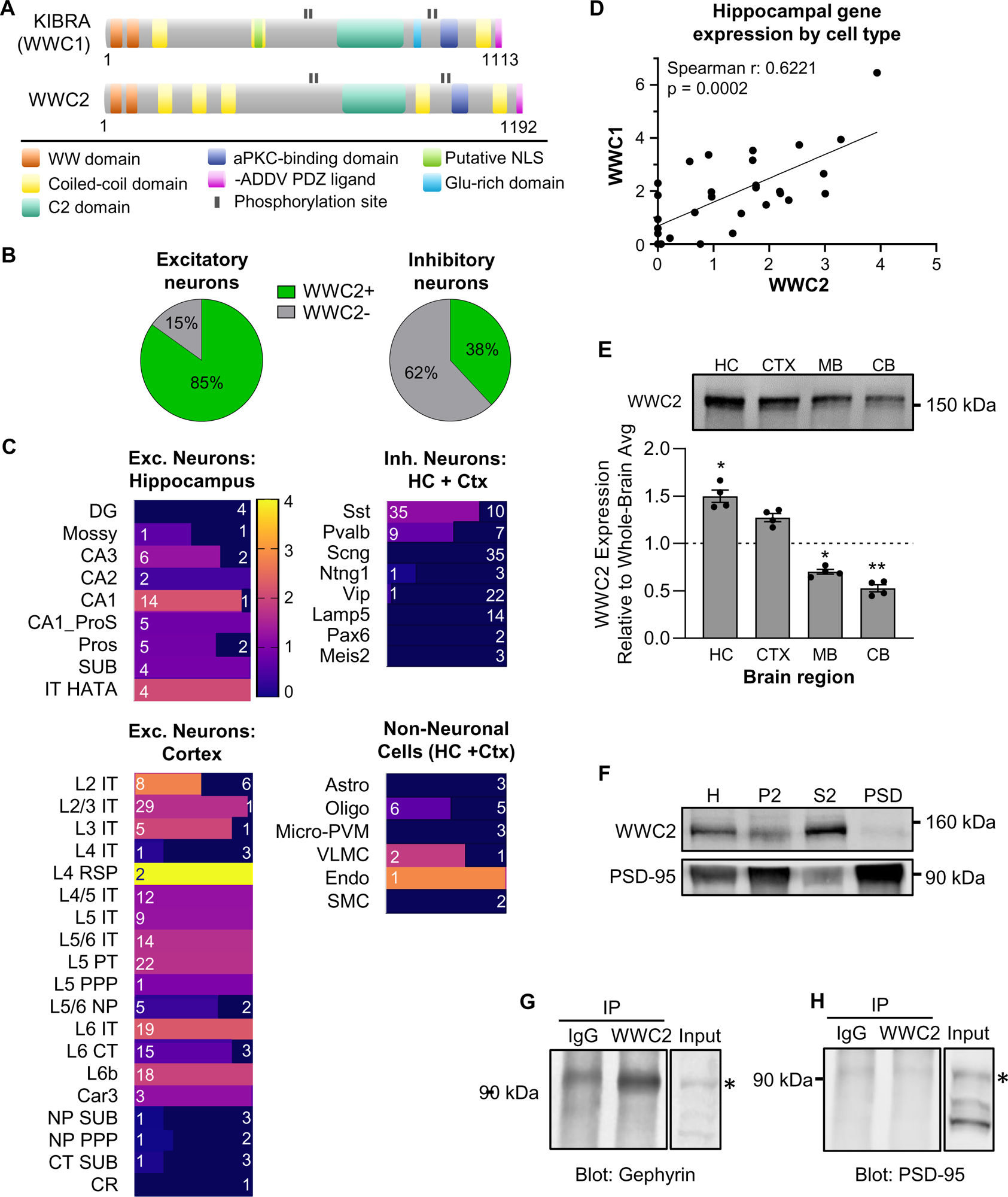
Distribution of WWC2 expression in the brain. **A)** Diagram of KIBRA and WWC2 interaction domains and phosphorylation sites. **B,C**) Analysis of WWC2 gene expression from Allen Institute for Brain Science cell types database: RNA-Seq Data, Mouse whole Cortex and Hippocampus-10x Genomics with 10X-Smart-Seq Taxonomy^32,83^. (B) Percent of identified excitatory (232) and inhibitory (121) neuron cell types from mouse hippocampus and cortex that express WWC2. (C) Number of each hippocampal excitatory, cortical excitatory, cortical + hippocampal inhibitory, and non-neuronal cell class that shows *Wwc2* expression. Dark blue portion of the bar represents the number of cells types in the indicated class that do not express *Wwc2*. Colored portion of the bar shows the number of cells types in the indicated class that express *Wwc2*, where the color represents the relative expression level (trimmed means). Numbers indicate number of cell types in each bar. For example, of the 45 somatostatin+ cell types, 35 express *Wwc2*, with relative expression of 1.17, averaged across all of the 35 Sst+ cell types that showed detectible *Wwc2* expression. **D)** Spearman correlation of WWC2 and WWC1 expression for each excitatory hippocampal cell type (15 CA1, 2 CA2, 8 CA3, 2 Mossy and 4 DG cell types). **E)** WWC2 is enriched in the hippocampus and cortex of WT animals. HC=hippocampus, CTX=cortex, MB=midbrain, CB=cerebellum. (***p=0.0006, repeated measures ANOVA. Multiple comparisons to average whole brain expression with Šidák correction: HC and MB *p < 0.05, CB **p < 0.01, CTX p = 0.0595.) N = 2 males + 2 females **F)** WWC2 is enriched in the cytosol (S2), present in the membrane-associated fraction (P2), and is depleted from the excitatory postsynaptic density (PSD). **G-H**) WWC2 interacts with gephyrin, but not PSD-95. Extra lanes removed for clarity.

γ-aminobutyric acid type-A receptors (GABA_A_Rs) are the primary mediators of fast ionotropic inhibition in the brain. Trafficking of GABA_A_Rs is tightly controlled, as dysregulated GABA_A_R expression can lead to deleterious downstream effects, including altered neuronal excitation/inhibition (E/I) balance and circuit dysfunction ^26,27^. Consequently, dysfunctions in GABA_A_R expression and trafficking have been implicated in numerous CNS pathologies such as childhood encephalopathic epilepsy, febrile seizures, ischemia, neurodegenerative disorders, and ASD, highlighting the importance of understanding the mechanisms through which GABA_A_R trafficking is regulated ^28–31^.

We report that WWC2 is a novel negative regulator of membrane GABA_A_R content. Loss of WWC2 results in increased surface GABA_A_R expression and dysregulated baseline inhibitory synaptic strength in hippocampal pyramidal neurons. We additionally identify WWC2 as a regulator of neuronal morphology, reporting that hippocampal neurons lacking WWC2 exhibit decreased dendritic complexity. These data represent evidence of divergent, class-specific synaptic functions of KIBRA and WWC2 within the hippocampus and contribute to our understanding of how WWC proteins influence neuronal function and cognition.

## RESULTS

### WWC2 is enriched in excitatory hippocampal and cortical neurons

Analysis of single cell RNAseq data from mouse hippocampus and cortex ^32^ reveals that, similar to KIBRA ^22^, WWC2 is expressed in the majority of excitatory cortical and hippocampal neuron cell types (85% of identified excitatory cell types, Fig. 1B) but only a subset of inhibitory neurons (38% of inhibitory cell types, Fig. 1B). Within the forebrain, *Wwc2* gene expression is highest in CA1 and cortical L2/3 and 4/5 neurons (Fig. 1C). Somatostatin and parvalbumin interneurons represent the vast majority of *Wwc2-* expressing inhibitory neurons (Fig. 1C). Substantial Wwc2 gene expression was also present in endothelial and vascular/leptomeningeal cells in the hippocampus and cortex (Fig. 1C), consistent with studies demonstrating a critical role for WWC2 in vascular development ^33,34^. Many, though not all, forebrain neurons co-express KIBRA and WWC2, and their gene expression is positively correlated in excitatory hippocampal cell types (Fig. 1D). WWC3 was not included in these analyses as it is not present in the mouse genome ^24^.

Consistent with RNA expression data, quantification of regional WWC2 protein expression revealed that WWC2 enriched in the hippocampus (Fig. 1E). Analysis of hippocampal WWC2 expression across development showed peak expression between postnatal days (P) 7-14, corresponding with the period of peak hippocampal synaptogenesis ^35^ (Fig. S1).

### WWC2 interacts with the inhibitory postsynaptic scaffold gephyrin

We examined WWC2 subcellular localization via serial fractionation in hippocampal tissue ^21,36^. WWC2 is present in the cytosol and plasma membrane/crude synaptosome fraction (P2), and is depleted from the purified excitatory PSD (Fig. 1F). Data from proximity ligation proteomics suggest WWC2 may be part of the inhibitory postsynaptic complex ^25^, in contrast to KIBRA (WWC1) which is enriched in the excitatory postsynaptic density ^19^. We examined WWC2 interaction with synaptic complexes by immunoprecipitating WWC2 from whole-brain homogenate and probing for interactions with core inhibitory and excitatory synaptic scaffolds gephyrin and PSD-95, respectively. We found that gephyrin co-immunoprecipitated with WWC2 whereas the excitatory synapse scaffold PSD-95 did not (Fig. 1G,H). Together with iPSD proteomics ^25^, these data strongly indicate that WWC2 is part of the inhibitory postsynaptic complex.

### WWC2 negatively regulates membrane GABA_A_R content

We sought to determine the possible role of WWC2 in regulating expression and localization of synaptic neurotransmitter receptors in the hippocampus. Pairings of mice with constitutive, heterozygous *Wwc2* deletion failed to produce offspring (Fig. S2), consistent with a recent report ^34^ indicating that constitutive loss of WWC2 is embryonically lethal. We therefore generated a forebrain excitatory neuron-driven knockout model by mating animals with LoxP sites flanking *Wwc2* exon 5 (*Wwc2 f/f)* to animals expressing cre recombinase under the *Emx1* promoter (Fig. S2). The resulting WWC2 cKO (*Wwc2 f/-; Emx1-cre^+^*) mice exhibited a near-total loss of WWC2 from the hippocampus (Fig. 2A), were viable, and born at expected ratios. Importantly, hippocampal KIBRA expression was unaltered in WWC2 cKO animals (Fig. 2A), indicating a lack of compensation following WWC2 deletion.

**Figure 2.**
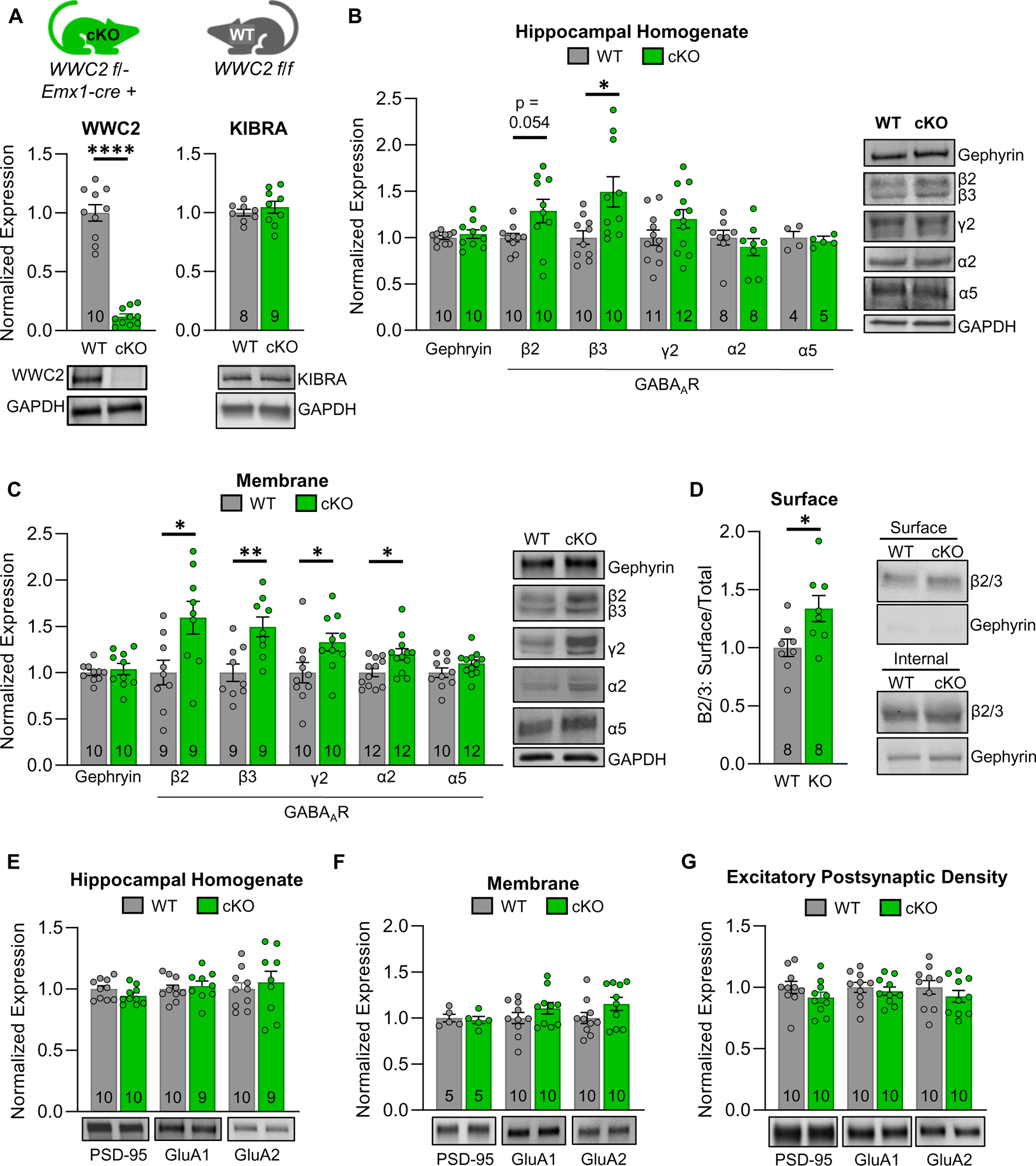
Increased surface GABA_A_R expression in WWC2 cKO hippocampus. **A)** Quantification of WWC proteins in *ex vivo* hippocampal homogenate from WT mice and WWC2 cKO littermates. (WWC2, WT = 1.0 ± 0.07, KO = 0.12 ± 0.02; KIBRA, WT = 1.0 ± 0.03, KO = 1.05 ± 0.05.Welch’s t-test p < 0.0001) **B,C**) Western blot analysis of gephyrin and GABA_A_R receptor subunits in *ex vivo* hippocampal homogenate (B) and membrane (C) fractions. (B, gephyrin, WT = 1.0 ± 0.02, KO = 1.04 ± 0.05; β2, WT = 1.0 ± 0.05, KO = 1.29 ± 0.13; β3, WT = 1.0 ± 0.07, KO = 1.49 ± 0.16; γ2, WT = 1.0 ± 0.08, KO = 1.20 ± 0.1; α2, WT = 1.0 ± 0.08, KO = 0.90 ± 0.09; α5, WT = 1.0 ± 0.07, KO = 0.96 ± 0.03. t-test with Welch’s correction *p=0.0163) (C, gephyrin, WT = 1.0 ± 0.03, KO = 1.04 ± 0.06; β2, WT = 1.0 ± 0.13, KO = 1.59 ± 0.18; β3, WT = 1.0 ± 0.09, KO = 1.49 ± 0.11; γ2, WT = 1.0 ± 0.11, KO = 1.33 ± 0.10; α2, WT = 1.0 ± 0.04, KO = 1.2 ± 0.06; α5, WT = 1.0 ± 0.05, KO = 1.1 ± 0.04. Welch’s t-test *p<0.05 [β2 p=0.0166, γ2 p=0.0373, α2 p=0.0156], **p=0.0032) **D**) Quantification of surface biotinylated proteins from acute hippocampal slices prepared from WWC2 cKO mice or WT littermates. Data presented as ratio of surface GABAR to total GABAR normalized to gephyrin signal. (WT = 1.0 ± 0.08, KO = 1.34 ± 0.11. Welch’s t-test *p=0.0247) **E-G**) Western blot analysis of PSD-95 and AMPAR subunits GluA1 and GluA2 in *ex vivo* hippocampal homogenate (E), membrane (F), and purified postsynaptic density (G) fractions. (E, PSD-95, WT = 1.0 ± 0.03, KO = 0.95 ± 0.27; GluA1, WT = 1.0 ± 0.03, KO = 1.03 ± 0.04; GluA2, WT = 1.0 ± 0.05, KO = 1.06 ± 0.09. F, PSD-95, WT = 1.0 ± 0.04, KO = 0.98 ± 0.04; GluA1, WT = 1.0 ± 0.06, KO = 1.11 ± 0.06; GluA2, WT = 1.0 ± 0.06, KO = 1.15 ± 0.07. G, PSD-95, WT = 1.0 ± 0.05, KO = 0.92 ± 0.04; GluA1, WT = 1.0 ± 0.04, KO = 0.97 ± 0.03; GluA2, WT = 1.0 ± 0.06, KO = 0.94 ± 0.05. No significant differences, Welch’s t-test). N, indicated in each bar, represents number of animals. Data presented as mean ± SEM. See Methods for sexes represented in each experimental group.

We fractionated hippocampal tissue from 1 month old (p28-35) WWC2 cKO animals and wild-type (WT, *Wwc2 f/f*) littermates, then analyzed ionotropic neurotransmitter receptor content in whole-cell, plasma membrane/crude synaptosome, and PSD fractions. GABA_A_R subunit β3 expression was significantly increased in cKO hippocampal homogenate, and β2 exhibited a trend toward increased expression (Fig. 2B). GABA_A_R subunits α2, β2, β3, and γ2 were all significantly increased in the membrane fraction of cKO animals (Fig. 2C). Thus, loss of WWC2 leads to aberrantly high GABA_A_R membrane expression, a finding corroborated by quantification of surface-expressed β2 and β3 via surface biotinylation of acute hippocampal slices (Fig. 2D). This upregulation appears to be specific for GABA_A_R subunits which are incorporated into the synapse, as expression of the predominantly extrasynaptic subunit α5 is unchanged in both the whole-cell and membrane fractions (Fig. 2B,C). Expression of total and membrane-associated gephryin were unchanged (Fig. 2B,C), indicating that the increase in GABA_A_R membrane expression was not secondary to increases in gephryin expression. Consistently, we do not observe changes in gephryin puncta size or density in cultured hippocampal neurons from WWC2 cKO mice (Fig. S3). Notably, expression of the AMPAR subunits GluA1 and GluA2 was unchanged in the whole-cell, membrane, and excitatory synapse (PSD) fractions (Fig. 2E-G) which contrasts with the depletion of extrasynaptic AMPARs observed upon KIBRA deletion ^21^. Expression of the excitatory synapse scaffold PSD-95 was also unchanged in hippocampal whole-cell, membrane, and PSD fractions, and PSD-95 puncta size and density were unaltered in cultured hippocampal neurons from WWC2 cKO mice (Fig. S3). Thus, loss of WWC2 produces substantial changes in membrane GABA_A_R expression, with no impact on AMPAR expression at excitatory synapses.

### GABA_A_R upregulation does not occur in KIBRA KO animals

KIBRA is well-established as a regulator of hippocampal AMPAR expression and trafficking ^19–21^. However, whether KIBRA also regulates expression or trafficking of GABA_A_Rs has yet to be established. To test whether the role of WWC2 as a regulator of GABA_A_R expression and localization is specific to WWC2 within the WWC family, we fractionated hippocampal tissue from KIBRA KO mice ^19^, then examined the expression of gephyrin and GABA_A_R subunits β2/3 and γ2. Total and membrane expression of gephyrin and GABA_A_R subunits was unaffected by KIBRA deletion (Fig. 3A,B). These data strongly suggest that KIBRA and WWC2 regulate the expression and/or trafficking of distinct receptor classes – with KIBRA regulating AMPARs (Fig. 3 and ^19–21^) and WWC2 regulating GABA_A_Rs (Fig. 2).

**Figure 3.**
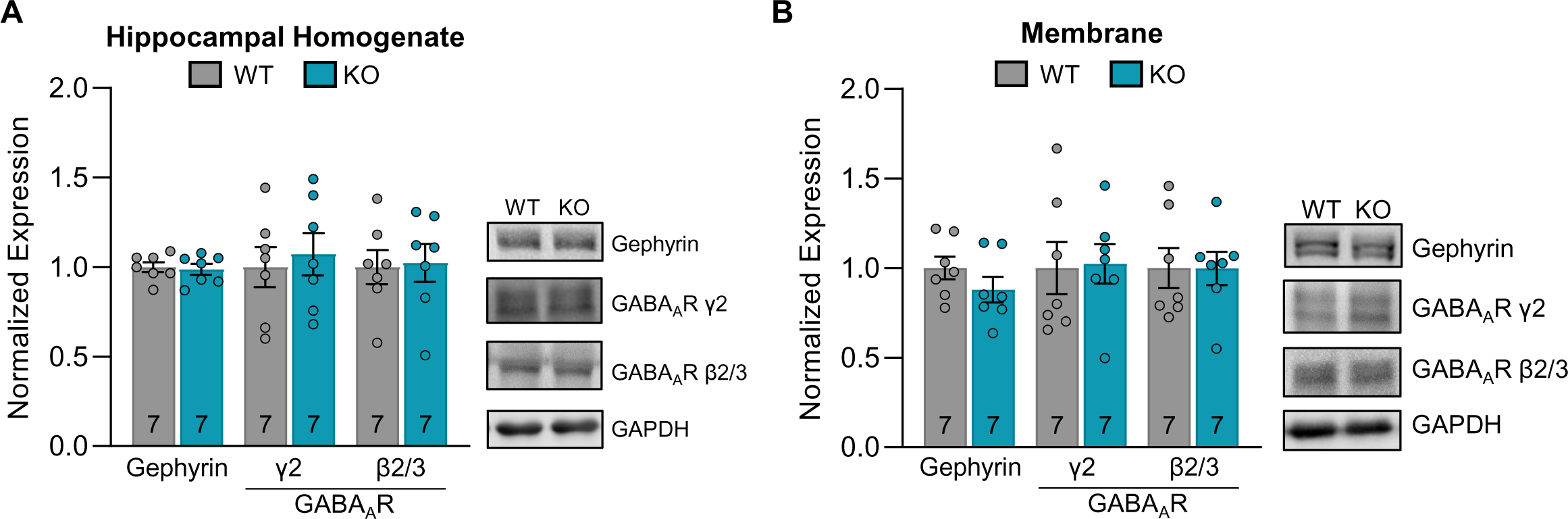
KIBRA loss does not affect GABA_A_R protein expression or membrane association. Western blot analysis of gephryin and GABA_A_R subunits γ2 and β2/3 protein expression in hippocampal homogenate (A) or membrane fraction (B) from WT mice and constitutive KIBRA KO littermates. (A, gephyrin, WT = 1.0 ± 0.03, KO = 0.99 ± 0.03; γ2, WT = 1.0 ± 0.11, KO = 1.07 ± 0.12; β2/3, WT = 1.0 ± 0.09, KO = 1.02 ± 0.11. B, gephyrin, WT = 1.0 ± 0.06, KO = 0.88 ± 0.07; γ2, WT = 1.0 ± 0.15, KO = 1.02 ± 0.11; β2/3, WT = 1.0 ± 0.11, KO = 1.0 ± 0.09.) N, indicated in each bar, represents number of animals (4 males + 3 females for each group). Data presented as mean ± SEM.

### The GABA_A_R recycling proteins HAP1 and GRIP1 are enriched in the membrane fraction of the WWC2 cKO hippocampus

GABA_A_R trafficking is a complex and highly dynamic network of processes encompassing plasma membrane insertion, clustering, removal, and intracellular recycling ^37–39^. Given that our data identify WWC2 as a negative regulator of cell surface GABA_A_R expression, we hypothesized that it participates in this process via one of three pathways: inhibition of GABA_A_R insertion or clustering, promotion of GABA_A_R internalization, or inhibition of endocytosed GABA_A_R recycling. We found no changes in expression or cleavage of the GABA_A_R insertion-promoting protein GABARAP (GABA_A_R-associated protein) ^40–42^ or in the phosphorylation of membrane γ2-S327 and β3-S408/409, which promote GABA_A_R synaptic clustering and prevent GABA_A_R endocytosis when phosphorylated, respectively (Fig. S4) ^43–46^. However, we found that huntingtin-associated protein 1 (HAP1) and glutamate receptor interacting protein 1 (GRIP1), which promote GABA_A_R recycling towards the plasma membrane ^47–50^, exhibit aberrantly high expression in the membrane fraction of the WWC2 cKO hippocampus (Fig 4A). The overabundance of membrane-associated GRIP1 in the WWC2 cKO is accompanied by a decrease in cytosolic GRIP1 (Fig. 4A). We also find that endogenous HAP1 and GRIP1 co-immunoprecipitate with endogenous WWC2 (Fig. 4B,C). Our findings therefore support the hypothesis that WWC2 interacts with the HAP1/GRIP1 complex to inhibit recycling of GABA_A_Rs back to the plasma membrane.

**Figure 4.**
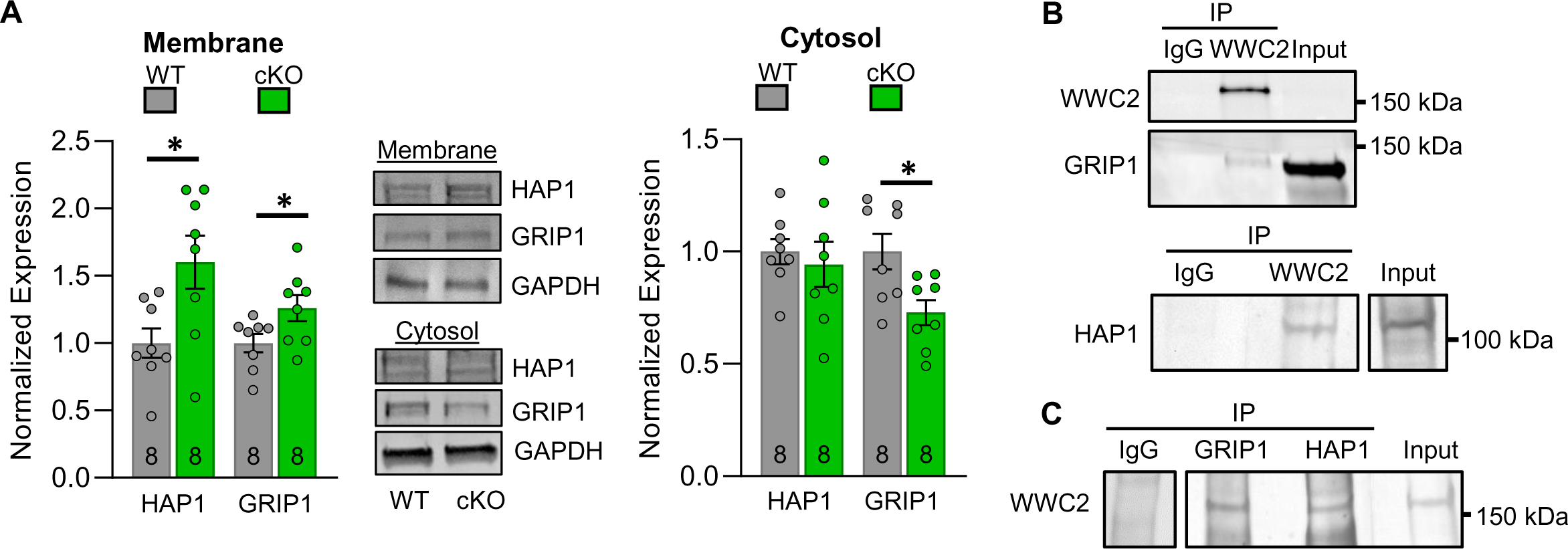
Loss of WWC2 increases membrane association of HAP1 and GRIP1. **A)** Quantification of HAP1 and GRIP1 protein expression in the membrane associated (left) and cytosolic (right) fractions from WWC2 cKO and WT hippocampal tissue. (Membrane: HAP1, WT = 1.0 ± 0.11, KO = 1.60 ± 0.20, *p = 0.0219; GRIP1, WT = 1.0 ± 0.07, KO = 1.26 ± 0.10, *p = 0.0483. Cytosol: HAP1, WT = 1.0 ± 0.06, KO = 0.94 ± 0.10; GRIP1, WT = 1.0 ± 0.08, KO = 0.73 ± 0.06, *p = 0.0150, Welch’s t-test.). N, indicated in each bar, represents number of animals (WT = 3 males + 5 females, KO = 4 males + 4 females). Data presented as mean ± SEM **B)** GRIP1 and HAP1 co-immunoprecipitate with WWC2 in forebrain homogenate from WT mice. **C)** WWC2 co-ommunoprecipitates with GRIP1 and HAP1 in forebrain homogenate from WT mice.

### Loss of WWC2 leads to dysregulation of inhibitory synaptic strength

To test the functional consequences of WWC2 loss, we induced sparse *Wwc2* deletion via biolistic transfection of *Wwc2 f/f* hippocampal slice cultures with mCherry-cre. Paired whole-cell recordings from neighboring cre-negative (WT) and cre-positive (cKO) CA1 pyramidal neurons revealed dysregulated miniature inhibitory postsynaptic current (mIPSCs) amplitude distribution in WWC2 cKO cells (Fig. 5A-C). Loss of WWC2 produced a bidirectional shift in mIPSC amplitudes with more large and more small mIPSCs (Fig. 5C), resulting in unchanged average mIPSC amplitude (Fig. 5D). To determine the cause of the distribution change, we examined the amplitude of cKO mIPSCs greater or less than 1 standard deviation from the WT mean. In WWC2 cKO neurons, mIPSC amplitude was significantly decreased for the small (<µ_WT_-σ) mIPSC population and significantly increased for the large (>µ_WT_+σ) mIPSC population (Fig. 5E). Further analysis indicated that the variability of mIPSC amplitude within individual WWC2 cKO cells was not different, rather cells segregated into two groups with largely unimodal shifts towards larger or smaller mIPSCs (Fig. S5). mIPSC kinetics were not different in cKO neurons (Fig. S5). These data indicate that WWC2 likely functions to normalize baseline inhibitory synaptic strength at the population level, consistent with a role in regulating GABA_A_R trafficking. WWC2 cKO neurons showed no change in mIPSC frequency (Fig. S5), consistent with our findings that the density of gephryin puncta is unchanged in hippocampal neurons cultured from *Wwc2* cKO mice (Fig. S3). Together, these data suggest that WWC2 regulates GABA_A_R trafficking without significantly contributing to the number of inhibitory postsynaptic sites.

**Figure 5.**
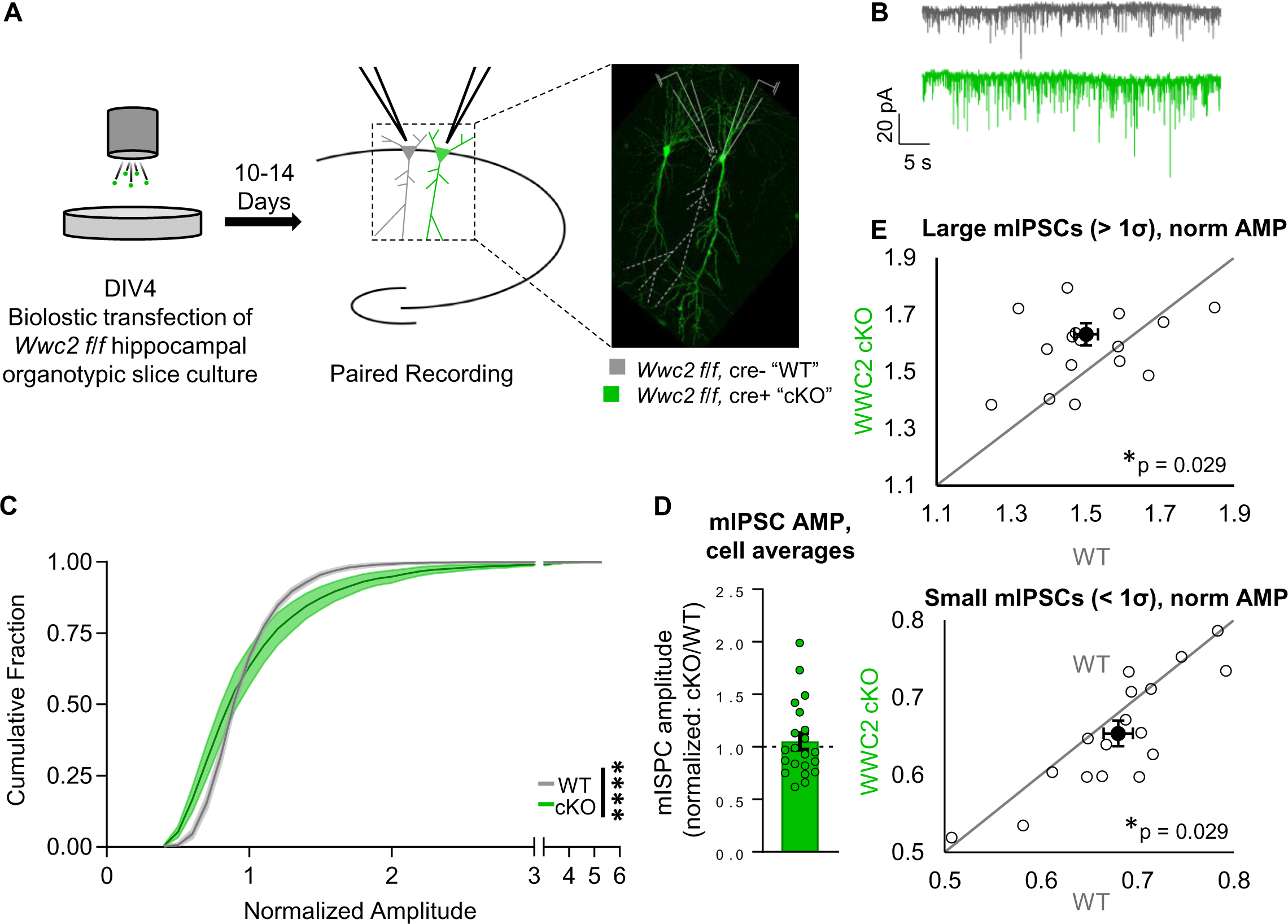
WWC2 cKO neurons exhibit dysregulated mIPSC amplitudes. **A)** Experimental schematic for paired recording of WT and WWC2 cKO neurons from biolistically-transfected organotypic hippocampal slice cultures. Note that color conventions carry throughout the figure. **B)** Sample mIPSC traces. **C)** Cumulative distribution of mIPSC amplitudes normalized to paired WT recording. N=21 paired recordings. Data presented as bin average ± SEM (shading) for bins sized 0.1.(**** p<0.0001, Kolmogorov-Smirnov test on unbinned data.) **D)** Average mIPSC amplitude across 21 recorded cKO cells, normalized to paired WT control. (cKO = 1.05 ± 0.08. 1-sample Wilcoxon test, not significantly different from 1.) **E)** Average normalized amplitude of mIPSCs greater or less than standard deviation from the WT mean. “Large” (>µWT+σ) mIPSCs are significantly larger and “small” (<µWT-σ) mIPSCs are significantly smaller in WWC2 cKO neurons. (Large WT=1.50 ± 0.03, cKO=1.63 ±0.04; Small WT=0.68 ± 0.02, cKO=0.65 ± 2. *p=0.029, paired t-test with Holm-Šídák correction for multiple comparisons.) C-E, n = 21 cell pairs (21 WT and 21 cKO) from 3 independent litters.1-2 cell pairs recorded per slice.

### WWC2 regulates dendritic arborization in hippocampal neurons

Loss of WWC2 also resulted in abnormal intrinsic membrane properties. WWC2 cKO neurons exhibited decreased capacitance (Fig. 6A), increased input resistance (IR) (Fig. 6B), and hyperpolarized resting membrane potential (Fig. 6C) compared to WT neurons. Changes in tonic inhibition mediated by extrasynaptic GABA_A_Rs can affect resting membrane potential^51^. However, membrane expression of α5 GABA_A_Rs, the predominant subunit mediating tonic inhibition in CA1 pyramidal neurons^52,53^, is unchanged in WWC2 cKO hippocampal tissue (Fig. 2C).

**Figure 6.**
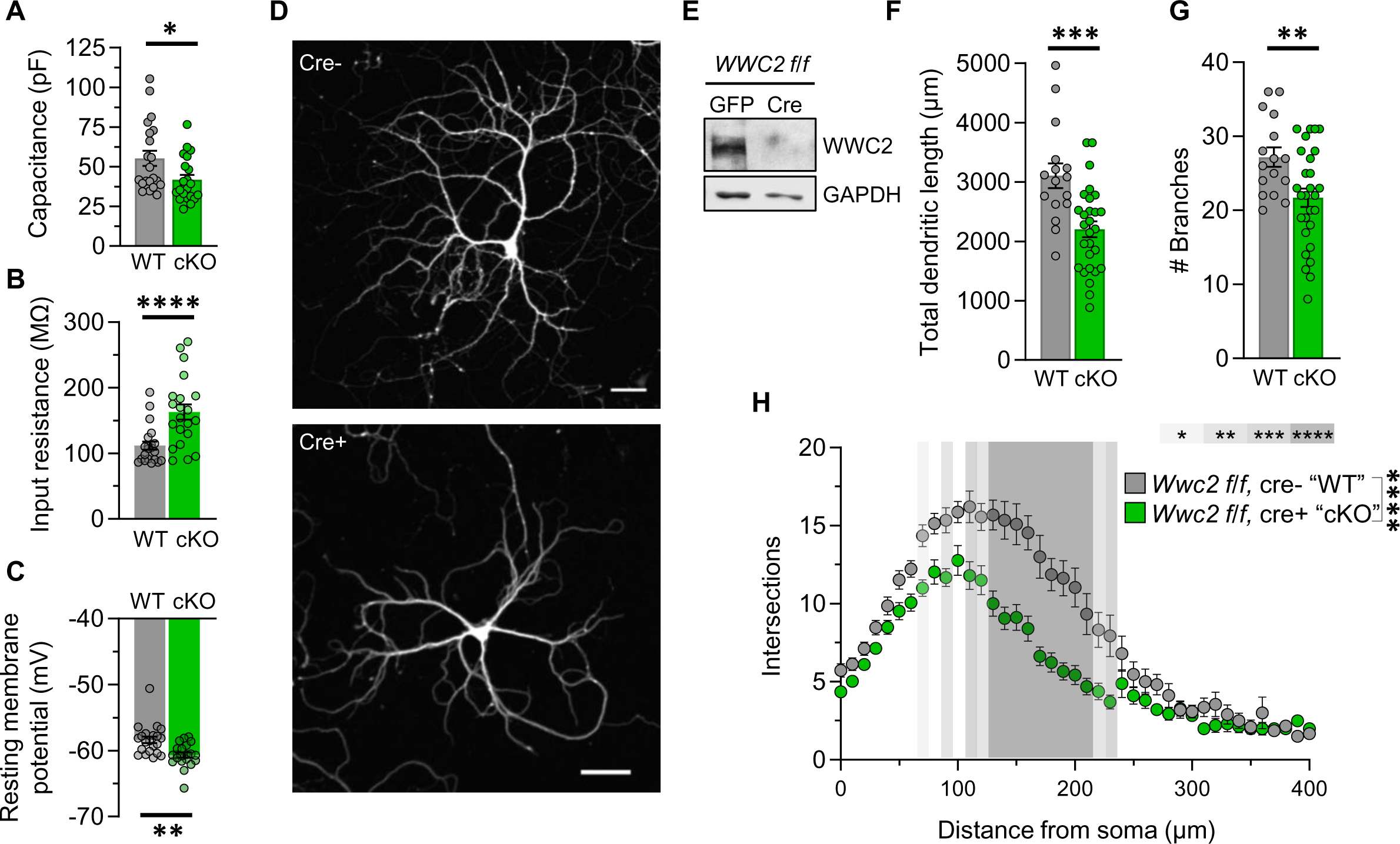
WWC2 loss results in decreased dendritic branching and aberrant intrinsic membrane properties. **A-C)** Intrinsic membrane properties from paired recording of WT and WWC2 cKO neurons in biolistically-transfected organotypic hippocampal slice cultures, as described in Figure 5. (A) WWC2 cKO neurons exhibit decreased capacitance. (WT = 55.2 ± 4.74, cKO = 41.7 ± 3.08. *p=0.014, paired t-test.) (B) Input resistance is increased in WWC2 cKO neurons. (WT = 111.7 ± 6.384, cKO = 162.9 ± 11.62. ****p<0.0001, paired Wilcoxon test.) (C) Resting membrane potential is hyperpolarized in WWC2 KO neurons. (WT = -58.38 ± 0.51, cKO = -60.65 ± 0.39. **p=0.0025, paired Wilcoxon test). **D)** Representative images of WT (cre-negative, top) and WWC2 cKO (cre-positive, bottom) cultured hippocampal neurons. Scale bars 50 µm. **E)** Western blot of WWC2 expression in cultured *Wwc2 f/f* hippocampal neurons receiving AAV-GFP (WT) or AAV-Cre (cKO). **F,G**) WWC2 cKO neurons have decreased total dendritic length (F) and number of branches (G) compared to WT controls. (F, WT = 3105 ± 209, KO = 2203 ± 135, **p=0.0011 Welch’s t-test. G, WT = 27.2 ± 1.31, KO = 21.71 ± 1.25, **p=0.0046 Welch’s t-test.) **G)** Sholl analysis of dendrites in cultured *Wwc2 f/f* neurons receiving AAV-GFP (WT) or AAV-GFP-cre (cKO). (2-way ANOVA: genotype p<0.0001; Šídák’s multiple comparisons post-hoc *p<0.05, **p<0.01, ***p<0.001, ****p<0.0001.) A-C. n = 21 cell pairs (21 WT and 21 cKO) from 3 independent litters.1-2 cell pairs recorded per slice. F-H, n=23 cells (WT) and 30 cells (cKO) from 2 independent cultures.

Changes in membrane capacitance and input resistance often reflect altered neuronal morphology. As KIBRA has been shown to positively regulate dendritic branching ^20^, we hypothesized that WWC2 regulates neuronal structure in addition to its impact on inhibitory synaptic function. Cultured hippocampal neurons from *Wwc2 f/f* mice were infected with AAV9-EGFP-cre (cKO) or AAV9-EGFP (WT) beginning on DIV 3. Analysis of dendritic branching morphology at DIV 18 revealed an overall decrease in dendritic length (Fig. 6F) and branch number (Fig. 6G) in WWC2 cKO neurons compared to WT controls. Sholl analysis further revealed deficits in dendritic complexity in WWC2 cKO neurons compared to WT controls (Fig. 6H). These finding are consistent with the decreased capacitance and increased IR seen in WWC2 cKO neurons (Fig. 6A,B). WWC2 cKO neurons in dissociated cultures and organotypic slice culture did not show a change in soma size (Fig. S6). Together, these data identify WWC2 as a regulator of both inhibitory synaptic function and dendritic morphology in hippocampal neurons.

## DISCUSSION

WWC family proteins are associated with several neurodevelopmental and neuropsychiatric disorders, highlighting a role in complex cognition ^9,10,13,14,16^. However, the cellular and molecular function of WWC2 in the brain was unknown prior to this study. We identify WWC2 as a component of the inhibitory postsynaptic complex that regulates GABA_A_ receptor membrane expression and inhibitory synaptic transmission. Additionally, we demonstrate a role for WWC2 in promoting dendritic branching, indicating that WWC2 regulates both neuronal structure and synapse function.

Strikingly, while early studies predicted similar or redundant cellular functions for WWC family proteins in the brain ^19,54^ our data demonstrate unique, synapse class-selective localization and function for WWC2 and KIBRA. WWC2 forms a complex with the inhibitory synapse scaffold gephryin and is present in plasma membrane fractions, whereas it is depleted from the excitatory postsynaptic density and does not interact with the core excitatory synapse scaffold PSD-95. Conversely, KIBRA is enriched in the excitatory postsynaptic density ^19^ and forms a complex with PSD-95 ^55^. Moreover, our data reveal a conserved role for WWC family proteins in synapse regulation through trafficking distinct classes of ionotropic neurotransmitter receptors; WWC2 regulates plasma membrane levels of GABA_A_Rs but not membrane or synaptic expression of AMPARs, whereas KIBRA regulates trafficking of AMPARs ^19–21^ but does not affect plasma membrane levels of GABA_A_Rs.

Our efforts to discern how WWC2 participates in regulating GABA_A_R trafficking revealed that deletion of WWC2 does not affect expression or cleavage of GABARAP or phosphorylation of membrane γ2-S327 and β3-S408/409, which drive GABA_A_R insertion, synaptic clustering, and surface retention, respectively ^41,42,46,56,57^. Rather, we demonstrate that WWC2 interacts with proteins that promote GABA_A_R recycling towards the plasma membrane, HAP1 and GRIP1 ^47–50,58^, and show that membrane expression of HAP1 and GRIP1 is increased in the WWC2 cKO hippocampus. These data are therefore consistent with a model in which WWC2 restricts recycling of endocytosed GABA_A_Rs back to the plasma membrane, although our data do not rule out a role for WWC2 in additional GABA_A_R trafficking steps.

As our data examined complexes of endogenous WWC2, GRIP1, and HAP1, it is not yet known whether WWC2 binds directly to GRIP1 and/or HAP1. A likely candidate for WWC2-GRIP1 interaction is through the c-terminal PDZ ligand of WWC2 interacting with one of the 7 PDZ domains in GRIP1. Additionally, HAP1 contains proline-rich sequences which may bind N-terminal WW domains in WWC2. Additional work will be needed to determine whether WWC2 binds GRIP1 and HAP1 individually or as a complex. Furthermore, whether WWC2 negatively regulates GABA_A_R recycling via interaction with GABA_A_R-bound HAP1/GRIP1 or through sequestering HAP1/GRIP1 away from GABA_A_Rs remains an open question. Interestingly, GRIP1 was originally identified as an AMPA receptor-binding protein ^59^ that promotes AMPAR trafficking and surface expression ^60,61^, and KIBRA was recently shown to interact with GRIP1 ^55^, suggesting that WWC proteins may utilize partially convergent mechanisms to regulate ionotropic neurotransmitter receptors at distinct classes of synapses.

Inhibitory GABA_A_R-mediated neurotransmission plays a critical role in regulating network stability and coordinating oscillations of neuronal ensembles that facilitate information transfer within and between brain regions ^62–64^. Dysregulation of mIPSC amplitude in WWC2 cKO neurons indicates that altered GABA_A_R trafficking in WWC2 cKO tissue has functional consequences. The broadening of the mIPSC amplitude distribution curve suggestes a failure of WWC2 cKO neurons to normalize the strength of their inhibitory synapses. Importantly, the variability of mIPSC amplitudes was unchanged within individual WWC2 cKO neurons; rather, WWC2 cKO neurons preferentially weaken or strengthen their inhibitory synapses as indicated by largely unimodal shifts in the mIPSC amplitude distribution in 18/21 cells pairs. The variability in inhibitory synaptic function is therefore present at a population level and likely has implications for circuit-level function. Aberrantly high rates of GABA_A_R recycling may lead to accumulation of unclustered, extrasynaptic GABA_A_Rs in the cells with smaller mIPSCs and accumulation of synaptic GABA_A_Rs in cells with larger mIPSCs. Whether the bidirectional, unimodal shifts in mIPSC amplitude indicate differential effects in two subpopulations of CA1 pyramidal cells (e.g. deep vs. superficial pyramidal neurons ^62,65^) is currently unknown.

While the precise mechanisms by which WWC2 and KIBRA complex with and regulate distinct classes of synapses remains to be determined, one intriguing hypothesis is that WWC2 and KIBRA facilitate assembly of membraneless inhibitory and excitatory postsynaptic signaling compartments (biomolecular condensates) via liquid-liquid phase separation (LLPS) ^66^. Both gephryin and PSD-95 complexes form biomolecular condensates in inhibitory and excitatory postsynaptic compartments, respectively ^67–69^. Additionally, KIBRA was recently shown to induce LLPS of Hippo signaling complexes in non-neuronal cells^70,71^. Both KIBRA and WWC2 contain multiple putative coiled-coil domains (2-3 in KIBRA, 5 in WWC2, UniProt 2023), which have been shown to mediate phase separation ^67,70,72–74^. Therefore, it is possible that that selective interaction with gephryin or PSD-95 and differences in coiled-coil domain content of WWC2 and KIBRA lead to preferential phase separation into the inhibitory and excitatory postsynaptic densities, respectively.

Interestingly, while the synaptic functions of WWC2 and KIBRA are distinct and class-specific (inhibitory vs. excitatory), the similarity between dendritic branching phenotypes in KIBRA knockdown ^20^ and WWC2 cKO neurons indicates that the two homologs may participate in similar mechanisms of neuronal morphology determination. As both WWC2 and KIBRA are known Hippo signaling activators, this may be via Hippo pathway components, which are expressed in post-mitotic neurons ^24,54,75^. Disrupted Hippo signaling has been associated with aberrant border cell protrusions and failure to maintain peripheral dendritic arborization in *Drosophila* ^76,77^. Loss of either KIBRA or WWC2 could decrease downstream LATS1/2 phosphorylation ^24,75^ and thereby interfere with dendrite polarization and the ability of neurons to grow or maintain their dendritic arbors. Future work should therefore consider the role of WWC proteins as Hippo regulators when investigating their roles in regulating neuronal structure.

The downstream circuit effects of altered inhibitory tone due to loss of WWC2 remain to be determined, but regulation of GABA_A_R expression and trafficking is of high functional importance for neural network function. Dysregulated E/I balance is implicated in many neurodevelopmental, psychiatric, and neurodegenerative disorders^78–80^. Of particular note is the increased GABA_A_R β3 subunit expression in WWC2 cKO tissue as *GABRB3* is implicated in seizure disorders^81,82^. Interestingly, gain-of-function GABRB3 mutations are associated with treatment resistance and more severe disease presentation than loss-of-function mutation ^82^.

In conclusion, we present the first data on the function of WWC2 in the CNS and identify divergent synaptic function within the WWC protein family. We demonstrate the necessity of WWC2 for normal GABA_A_R trafficking, baseline synaptic strength, and dendritic morphology in hippocampal neurons, providing insight into how WWC proteins influence neuronal function and cognition via regulation of receptor trafficking and neuronal morphology.

## Acknowledgments

### Funding

National Institutes of Health Grant NIMH 1R01MH117149-01 (LJV)

### Author contributions

Conceptualization: LJV, TLD

Methodology: LJV, TLD, JRW

Resources: TLD, RCJ, RLH

Investigation: TLD, JRW

Formal Analysis: LJV, TLD

Visualization: LJV, TLD

Supervision: LJV

Writing—original draft: LJV, TLD

Writing—review & editing: LJV, TLD

## MATERIALS AND METHODS

### Experimental Model and Study Participant Details

The University of Texas Southwestern Institutional Animal Care and Use Committees approved all animal protocols in this study. Mice were group housed in a climate-controlled environment on a 12-hour light/dark cycle. Food and water were provided *ad libitum*. Hemizygous Emx1-cre mice^84^ (Jackson Labs strain # 005628,) were bred with homozygous *Wwc2 floxed/floxed* mice (C57Bl/6N, N10+, see Figure S2), yielding cre positive (cre+) *Wwc2 floxed/null* and cre negative (cre-) *Wwc2 floxed/floxed* experimental littermates. WWC2 KO animals were derived using a floxed over null approach, as preventing spontaneous recombination proved impossible when using cre under either the *Emx1* or *Syn1* promoters. Generation of KIBRA KO mice was described previously^19^ (C57Bl/6N, N10+). Both male and female mice were used in all experiments. See table below for details.

| Fig. 1 |  | WT |  |
| --- | --- | --- | --- |
| E | Protein | Males | Females |
|  | WWC2 | 2 | 2 |

| Fig. 2 |  | WT |  | KO |  |
| --- | --- | --- | --- | --- | --- |
|  | Protein | Males | Females | Males | Females |
| Homogenate | WWC2 | 6 | 4 | 6 | 4 |
|  | KIBRA | 3 | 5 | 4 | 5 |
|  | Gephyrin | 6 | 4 | 6 | 4 |
| | β2 | 3 | 7 | 3 | 7 |
| | β3 | 3 | 7 | 3 | 7 |
| | γ2 | 3 | 8 | 6 | 6 |
| | α2 | 4 | 4 | 3 | 5 |
| | α5 | 2 | 2 | 3 | 2 |
|  | PSD-95 | 6 | 4 | 6 | 4 |
|  | GluA1 | 3 | 7 | 3 | 6 |
|  | GluA2 | 3 | 7 | 3 | 6 |
| P2 | Gephyrin | 6 | 4 | 6 | 4 |
| | β2 | 3 | 6 | 3 | 6 |
| | β3 | 3 | 6 | 3 | 6 |
| | γ2 | 3 | 7 | 5 | 5 |
| | α2 | 5 | 7 | 5 | 7 |
| | α5 | 4 | 6 | 6 | 6 |
|  | PSD-95 | 3 | 2 | 3 | 2 |
|  | GluA1 | 3 | 7 | 3 | 7 |
|  | GluA2 | 3 | 7 | 3 | 7 |
| PSD | PSD-95 | 3 | 7 | 3 | 7 |
|  | GluA1 | 3 | 7 | 3 | 7 |
|  | GluA2 | 3 | 7 | 3 | 7 |
| Surface Biotin | β2/3 | 4 | 4 | 4 | 4 |

| Fig. 3 |  | WT |  | KO |  |
| --- | --- | --- | --- | --- | --- |
|  | Protein | Males | Females | Males | Females |
| Homogenate | Gephyrin | 4 | 3 | 4 | 3 |
| | β2/3 | 4 | 3 | 4 | 3 |
| | γ2 | 4 | 3 | 4 | 3 |
| P2 | Gephyrin | 4 | 3 | 4 | 3 |
| | β2/3 | 4 | 3 | 4 | 3 |
| | γ2 | 4 | 3 | 4 | 3 |

| Fig. 4 |  | WT |  | KO |  |
| --- | --- | --- | --- | --- | --- |
|  | Protein | Males | Females | Males | Females |
| P2 | HAP1 | 3 | 5 | 4 | 4 |
|  | GRIP1 | 3 | 5 | 4 | 4 |
| Cytosol | HAP1 | 3 | 5 | 4 | 4 |
|  | GRIP1 | 3 | 5 | 4 | 4 |

| Supp. Fig. 1 |  | WT |  |
| --- | --- | --- | --- |
|  | Protein | Males | Females |
|  | WWC2 | 2 | 2 |

| Supp. Fig. 4 |  | WT |  | KO |  |
| --- | --- | --- | --- | --- | --- |
|  | Protein | Males | Females | Males | Females |
| A | γ2-pS327 | 3 | 5 | 4 | 4 |
| | β3-pS327 | 3 | 5 | 4 | 4 |
| B | GABARAP | 4 | 5 | 4 | 6 |

### Method Details

#### Genotyping and primers

Genotyping for *Emx1-cre* and *Wwc2* floxed, WT, and KO alleles was performed via step-down polymerase chain reaction (PCR) amplification of DNA from tail biopsy tissue. *Emx1-cre* primers were: forward 5’-CAACGGGGAGGACATTGA-3’ and reverse 5’-TCGATAAGCCAGGGG TTC-3’ (Integrated DNA Technologies). *WWC2* floxed primers were: forward 5’-GGATGAACCAAATATCTTTCTGCCTC-3’ and reverse 5’-CTTAGTCTGGAAAATGGGGACCACG-3’. *WWC2* KO primers were: WT forward 5’-AAAAATAGGCGTATCACGAGGC-3’, KO forward 5’-GTGAATGTTGAGTAGAATCCCACTGC-3’, and floxed reverse 5’-CTTAGTCTGGAAAATGGGGACCACG-3’. 10 cycles of decreasing annealing temperatures (from 65-60 °C) followed an initial denaturation at 94 °C. Amplification then proceeded for 28 cycles, with an annealing temperature of 56 °C. PCR products were visualized on 1.5% agarose gels containing SyberSafe or GreenGlo dye. The following bands were detected: *Emx1-cre* mutant 195 bp, *Wwc2* WT 189 bp (floxed reaction) or 329 (KO reaction), *Wwc2* floxed 406 bp (floxed reaction) or 500 bp (KO reaction), and *Wwc2* KO 208 bp.

#### Brain Dissection

Prior to rapid decapitation, mice were anesthetized via cryoanesthesia if 0-1 day old or by exposure to isoflurane before being rapidly decapitated. The brain was placed in ice cold dissection buffer (in mM: 125 NaCl, 3.25 KCl, 25 NaHCO3, 1.25 NaH2PO4·H2O, 11 glucose, 0.75 CaCl2, 7 MgCl2) and the areas of interest isolated and pooled. For experiments using whole-brain homogenate, one hemisphere was left intact. Immediately following dissection, the tissue was flash frozen in liquid nitrogen and stored at -80 °C until processed for experimental use.

#### Tissue Preparation for Developmental & Regional WWC2 Expression Analysis

Regions of interest were homogenized in 500 µL (hippocampus) or 1mL (cortex, midbrain, cerebellum, whole brain) of RIPA buffer (for developmental analysis, 1% Triton X-100, 0.5% sodium deoxycholate, 0.1% SDS, 50 mM Tris-HCl, 100 mM NaCl, 1 mM EDTA, 50 mM NaF, 10 mM NaPPi, 1 mM NAVO3, and HALT protease inhibitors) or PBS with 1% Triton X-100 and HALT protease inhibitors (for regional analysis). Tissue was homogenized by repeated passage through a 26-gauge needle and syringe. Protein concentration was determined by using the Pierce^TM^ Detergent Compatible Bradford Assay Kit and a BioTek Synergy H1 Microplate Reader. Samples were boiled for 10 minutes in SDS protein sample buffer containing the following: 10% glycerol, 62.5mM Tris/HCl pH 6.8, 2% sodium dodecyl sulfate (SDS), 0.01% bromophenol blue, 1.25% beta-mercaptoethanol.

#### Immunoprecipitation

Forebrain tissue (both hemispheres from one mouse) was homogenized in 1 mL of immunoprecipitation buffer (PBS containing in mM: 150 NaCl and 1 EDTA with 1% Triton-X100 and HALT Protease Inhibitors). Protein concentration was determined by using the Pierce ^TM^ Detergent Compatible Bradford Assay Kit. 300 µg of homogenized forebrain was pre-cleared by centrifugation at 800xg for 10 minutes followed by a 1-hour incubation with Protein A/G magnetic beads. Following removal of the beads, 3 µg of anti-WWC2 antibody was added to the homogenized tissue and incubated overnight with rotation at 4°. Protein A/G magnetic beads were added to the primary antibody mixture and incubated for 1 hour at 4°C. Beads were collected by application of a magnetic field and washed three times in immunoprecipitation buffer and two times in PBS. Bound sample was eluted by incubation in SDS protein sample buffer for 10 minutes at room temperature.

#### Subcellular Fractionation

Pooled hippocampal tissue was homogenized in 1 mL of homogenization buffer containing the following (in mM): 320 sucrose, 10 HEPES at pH 7.4, 1 EDTA, 1 NaPPi, 5 NaF, 1 NaVO3, HALT Protease Inhibitor. 70 microliters of homogenate were removed for analysis of total protein expression. The remaining homogenate was then centrifuged at 800 xg for 10 minutes at 4°C. The supernatant (S1) was transferred to VWR high G-force micro-centrifuge tubes and centrifuged at 15,000 xg for 20 minutes at 4°C. The resultant supernatant (S2) was removed from the pellet (P2, membrane fraction), and the pellet was lysed in 500 microliters of Milli-Q water with protease and phosphatase inhibitors containing the following (in mM): 1 NaPPi, 5 NaF, 1 NaVO3, HALT Protease Inhibitor. To ensure complete lysis of P2, 2 microliters of 1M HEPES at pH 7.4 was added to each sample followed by incubation with agitation for 30 minutes at 4°C. 70 microliters were removed from each sample following incubation and stored for future analysis of the membrane fraction. The remaining lysed P2 fraction was then centrifuged at 25,000 xg for 20 minutes at 4°C. The lysed supernatant (LS1) was removed, and the lysed pellet (LP1) was resuspended in 250 mL of 50mM HEPES pH 7.4 with protease and phosphatase inhibitors. Each sample was then rapidly mixed with an equal volume of 250μL of 1% triton X-100 with protease and phosphatase inhibitors and incubated with agitation at 4°C for 15 minutes. The resuspended LP1 was then centrifuged at 32,000 xg for 20 minutes at 4°C. The lysed supernatant (LS2) was discarded, thereby leaving the lysed pellet or PSD fraction. The PSD fraction was resuspended in 100μL of 50 mM HEPES pH 7.4 with proteasome and phosphatase inhibitors. Protein concentrations of each fraction were determined using the Pierce^TM^ Detergent Compatible Bradford Assay Kit. Samples were boiled for 10 minutes in SDS protein sample buffer containing the following: 10% glycerol, 62.5mM Tris/HCl pH 6.8, 2% sodium dodecyl sulfate (SDS), 0.01% bromophenol blue, 1.25% beta-mercaptoethanol.

#### Surface Biotinylation

380 µm acute hippocampal slices were made from 28 to 32-day-old animals. Slices recovered at 30°C in oxygenated artificial CSF (ACSF) containing (in mM): 125 NaCl, 5 KCl, 1.25 NaH2PO4, 26 NaHCO3, 10 d-glucose, 2CaCl2, and 1 MgCl2. After recovery, slices were cooled with ice-cold ACSF then incubated with ACSF containing 1 mg/mL Sulfo-NHS-LC-Biotin for 25 minutes on ice. Excess biotinylation reagent was quenched by two twenty-minute washes in ACSF containing 100 mM glycine, on ice. Slices were washed again with ice-cold ACSF and homogenized via 30-minute incubation in RIPA buffer before mechanical dissociation through a 27g needle. The homogenates were centrifuged at 15,000 x g for 15 minutes at 4°C. 50 µg of supernatant was incubated with 25 µL of streptavidin-conjugated agarose beads overnight at 4°C. The agarose beads were washed with RIPA and the bound fraction eluted by incubation in 2x SDS-PAGE sample buffer for 30 minutes at room temperature before analysis of biotinylated proteins via immunoblotting.

#### Immunoblotting

Samples were loaded into either a 10% SDS-PAGE gel or 4-20% gradient SDS-PAGE gel. Following separation, the gels underwent a wet protein transfer to a Nitrocellulose membrane (Odyssey Nitrocellulose Membrane, pore size 0.22 μm) run in ice-cold transfer buffer at 100 V for 2 hours. The membranes were then incubated in ThermoScientific StartingBlock (TBS) for 20 minutes. Following blocking, the membranes were incubated in primary antibody diluted in TBS + 0.1% Tween-20 + 5% StartingBlock at 4°C overnight. Membranes were incubated with fluorescent secondary antibody for 1 hour and imaged on a Bio-Rad ChemiDoc MP. Primary antibodies were used at the following dilutions: 1:10,000 (GAPDH), 1:5,000 (PSD-95), 1:500 (KIBRA, GABA_A_R β2/3, GABA_A_R γ2-pS3273), 1:1,000 (all other antibodies). See Key Resources Table for additional information about individual antibodies.

#### Western Blot Analysis

Developmental expression data were quantified using the Li-Cor Image Studio Lite software. Samples were normalized to GAPDH as a loading control before being normalized to the within-litter P0 sample.

Fractionation data were quantified using the Li-Cor Image Studio Lite software. Homogenate, cytosolic, and membrane fraction samples were normalized to GAPDH and PSD samples were normalized to total protein. Total protein signals were visualized with No-Stain Protein Labeling Reagent (ThermoFisher, #A44449) according to manufacturer’s protocol and quantified in ImageJ. Loading control normalized signals were subsequently normalized to the average of all WT (WWC2*floxed/floxed*) littermate samples within the same gel. Note that GAPDH is present (though not enriched) in all fractions analyzed ^85^.

Surface biotinylation data were quantified using the Li-Cor Image Studio Lite software. Total β2/3 was normalized to gephyrin, then used to normalize the corresponding surface signal. Signals were then normalized to the average of all WT samples within the gel. Note that our data show that gephyrin membrane expression is unchanged in WWC2 cKO samples.

Quantification of phosphorylation was accomplished by first normalizing the total protein to the GAPDH loading control, then normalizing the phosphorylation signal to the normalized total signal. Normalized phosphorylation signal was then normalized to the average of WT samples within the gel.

#### Hippocampal slice culture and transfection

Organotypic hippocampal slice cultures were prepared from p6 pups as described in ^86^. Briefly, hippocampi were dissected in ice-cold dissection buffer (10 mM HEPES, 1mM CaCl_2_, 5mM MgCl_2_, 10 mM dextrose, 4mM KCl, 26 mM NAHCO_3_, 248 mM sucrose, pH 7.28) and sliced at a thickness of 400 µm using a tissue chopper. 6-8 slices per animal were transferred to cell culture inserts (Millipore #PICM 030 50) and maintained in tissue culture media (Minimal Essential Media + 1 mM L-glutamine, 0.0012% Ascorbic Acid, 1 mM CaCl_2_, 2 mM MgSO_4_, 12.87 mM dextrose, 5.25 mM NAHCO_3_, 30 mM HEPES, 0.17 mM insulin, pH 7.28, 310-315 mOsm) at 35 °C, 5% CO_2_. Slice cultures were biolistically transfected at 4 days *in vitro* (DIV). Biolistic transfection and bullet preparation were performed with the BioRad Helios Gene Gun system as described in^86,87^. mCherry-NLS-Cre and PA1-GFP constructs have been described previously^88,89^. Experiments were conducted 10 – 14 days post-transfection.

All studies utilized three independent slice cultures (litters). The experimental design for electrophysiology and imaging in organotypic hippocampal slice culture utilizes two sister culture inserts prepared from one animal (∼7-8 hippocampal slices per insert). For electrophysiology, all inserts were transfected with mCherry-NLS-Cre plus PA1-GFP. For imaging, slice culture inserts from the same animal were transfected with either mCherry-NLS-Cre plus PA1-GFP or vector plus PA1-GFP. Images were collected from sister inserts on the same day. For all studies, analysis was conducted by an investigator blind to transfection construct.

#### Electrophysiology

Simultaneous or sequential paired whole-cell voltage-clamp recordings were obtained from transfected (WWC2 knockdown) and neighboring untransfected (WWC2 wildtype) neurons under visual guidance using IR-DIC and GFP/mCherry fluorescence to identify transfected neurons as described ^86^. Transfected neurons were required to have GFP fluorescence and clear nuclear mCherry expression. Recordings from slice cultures were made at 28-30°C in a submersion chamber perfused at 3 ml/min with artificial cerebrospinal fluid (aCSF) containing (in mM): 119 NaCl, 2.5 KCl, 26 NaHCO_3_, 1 NaH_2_PO_4_, 11 D-Glucose, 3 CaCl_2_, 2 MgCl_2_; pH 7.28, 300 mOsm and saturated with 95%O_2_/5%CO_2_. For mIPSC measurements, aCSF was supplemented with 10μM CPP and 20μM DNQX and whole cell recording pipettes (∼4-7.5 MΩ) were filled with a high chloride intracellular solution containing (in mM): 0.2 EGTA, 79 K-gluconate, 44 KCl, 6 NaCl, 10 HEPES, 4 ATP-Mg, 0.4 GTP-Na, 14 phosphocreatine-Tris; pH 7.2, 290 mOsm. All chemical reagents were acquired from Millipore-Sigma, HelloBio, or Tocris. Iinput and series resistances were measured in voltage clamp with a 400-ms, –10 mV step from a –60 mV holding potential (filtered at 30 kHz, sampled at 50 kHz). Cells were only used for analysis if the following criteria were met: series resistance < 25 MΩ and stable throughout the experiment, resting membrane potential < -35mV; and input resistance > 75 MΩ. Waveforms were filtered at 3 kHz, acquired and digitized at 10 kHz on a PC using custom software (LabView; National Instruments).

Data analysis was performed using custom software in LabView ^86^. mEPSCs and mIPSCs were detected off-line using an automatic detection program (Clampfit; Molecular Devices, San Jose, CA.), with mIPSC detection based on a cell-specific averaged event template based on >5 events, followed by subsequent visual confirmation. The template match threshold remained constant for the duration of each experiment. For each cell pair, individual mIPSC measurements were normalized to the average value across all mIPSCs in the WT cell. mIPSCs were categorized as “small” or “large” if the amplitude fell outside of 1 standard deviation from the WT cell mean (small if <µ_WT_-σ, large if >µ_WT_+σ).

#### Hippocampal neuron culture

Hippocampi were dissected from P0-P1 mouse pups and incubated with papain and DNase at 37° for 15 minutes. Tissue dissociation was stopped by 2 washes with warmed neuronal plating medium (Neurobasal Medium supplemented with 5% horse serum, B-27 neuronal supplement, glutamax, and penicillin/streptomycin). The tissue was serially triturated using 3 fire-polished glass Pasteur pipettes with decreasing tip diameter and filtered through a 70µm cell filter to obtain dissociated neurons. The cells were pelleted by centrifugation for 4 minutes at 120xg and resuspended in fresh neuronal plating medium. Cell counting was performed using a hemocytometer. Neurons were plated on acid-etched glass coverslips coated for 1 hour in poly-L lysine at a density of 50,000 cells/mL for branching analysis or 100,000 cells/mL for synaptic puncta quantification. On DIV 1 and 3, half of the media was changed for serum-free neuronal maintenance medium (Neurobasal Medium supplemented with B-27 neuronal supplement, glutamax, and penicillin/streptomycin). Cultured cells were maintained at 37°C and in a 5% CO_2_ environment until fixation.

#### Dendritic branching analysis

Cultured neurons from WWC2*floxed/floxed* pups were infected with AAV9-EGFP-cre ((pENN.AAV9.hSyn.Cre.WPRE.hGH, Addgene #105553-AAV9) or AAV9-GFP (pENN.AAV.CB7.CI.eGFP.WPRE.rBG, Addgene #105542-AAV9) on DIV 3. Cells were then reinfected on DIV 10 with AAV9-mCherry (pENN.AAV9.CB7.CI.mCherry.WPRE.RBG, Addgene #105544-AAV9). Cells were fixed on DIV 18 and endogenous mCherry fluorescence was imaged using a Nikon C2 scanning confocal microscope. Single z-plane images containing the entire dendritic tree were acquired with a 20x air objective. Images were analyzed with the ImageJ plugin SNT. Cells plated in poly-L lysine coated 60 mm plates and treated with the aforementioned viruses were scraped and homogenized in RIPA buffer on DIV 18. WWC2 knockout was confirmed via immunoblotting of these samples.

#### Immunofluorescence

Cultured neurons were plated in separate wells. Cells were fixed at DIV 17-19 as previously described and permeabilized via 10-minute incubation in 0.2% Triton-X100 in PBS. Nonspecific binding sites were blocked by incubation in antibody buffer (containing, in mM: 150 NaCl, 50 Tris-Base, 100 L-lysine, and 1% bovine serum albumin) supplemented with 50% normal goat serum (NGS) for one at room temperature. Coverslips were incubated with primary antibodies diluted in antibody buffer + 10% NGS overnight at 4° with shaking. Following washing with PBS, coverslips were incubated with fluorescent secondary antibodies diluted in antibody buffer + 10% NGS for 2 hours, in the dark, at room temperature. Coverslips were mounted using FluoroShield and sealed with clear nail polish.

#### Synapse quantification

Images of synaptic puncta were acquired on a Zeiss LSM880 Airyscan laser scanning confocal microscope using a 63x oil-immersion objective and a 1.8x optical zoom. Z-stacks were taken at a resolution of 2048×2048 at 0.185-micron intervals through the entirety of the region of interest, then subjected to 3D Airyscan deconvolution. Maximum intensity projections of the resultant z-stacks were used for analysis in ImageJ. Image thresholds were set at +2 standard deviations from the mean fluorescence in each channel and the resultant puncta were analyzed using Particle Analyzer in ImageJ. Regions of interest were manually drawn to include either the first ∼100 µm of primary dendrite or the soma. Synapse density was quantified by dividing the number of puncta by the length of dendrite or area of soma analyzed. Synapse area data are presented as the average synapse size per ROI, averaged across all ROIs quantified. Only one dendritic and somatic ROI were analyzed for each neuron.

#### Quantification and statistical analysis

Statistical analysis was performed in Graph Pad Prism version 9.4.1. Data are plotted as mean ± SEM unless otherwise noted. The statistical test performed for each analysis is noted in the figure legend. p< 0.05 was considered significant. Data and test residuals were assessed for normality using Shapiro-wilk and D’Agostino-Pearson tests, in addition to visual inspection of Q-Q plots. Nonparametric tests (Wilcoxon, Mann-Whitney, Kolmogorov-Smirnov) were used in the small number of cases where normality criteria were not met. Equality of variance between groups was assessed using an F test (t-tests) and examination of residual homoscedasticity plots. For data that did not meet criteria for equal variance a Welch’s correction was applied to the statistical test. Sphericity (equal variability of differences) was not assumed for repeated measures ANOVA (Geisser-Greenhouse correction was applied).The Šidák correction was applied to ANOVA post-hoc multiple comparisons. mIPSC cumulative distributions were compared using the Kolmogorov-Smirnov test on unbinned data. Statistical tests were chosen based on sample size, hypotheses, and agreement with statistical assumptions.

---

**Figure S1.**
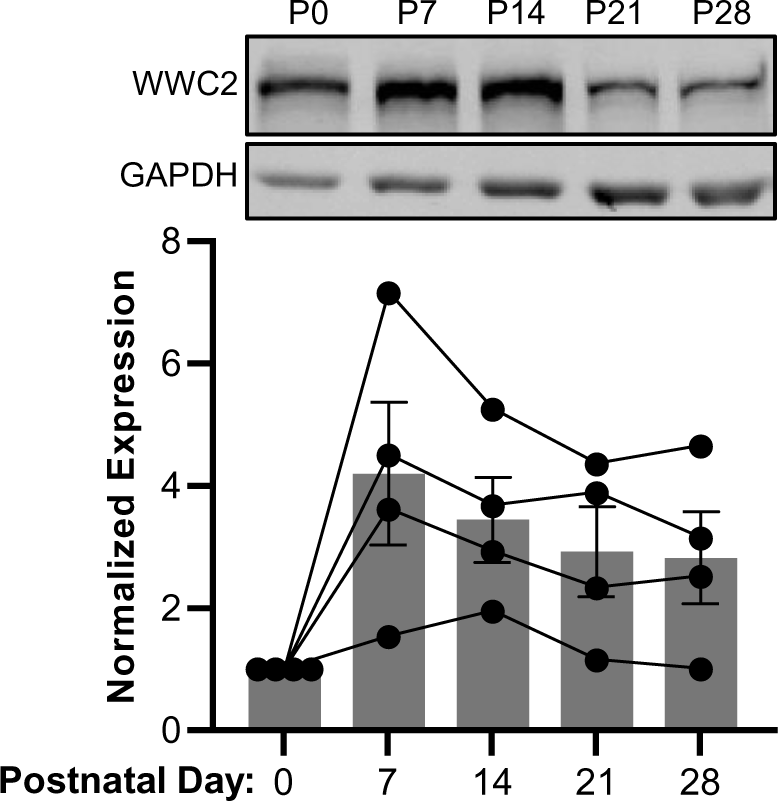
Developmental expression of WWC2. WWWC2 protein expression was assessed by western blot from hippocampal homogenate prepared from mice at P0, 7, 14, 21, and 28. Lines indicate mice from the same litter. N = 4 mice per time point (2 males + 2 females). Data presented as mean ± SEM

**Figure S2.**
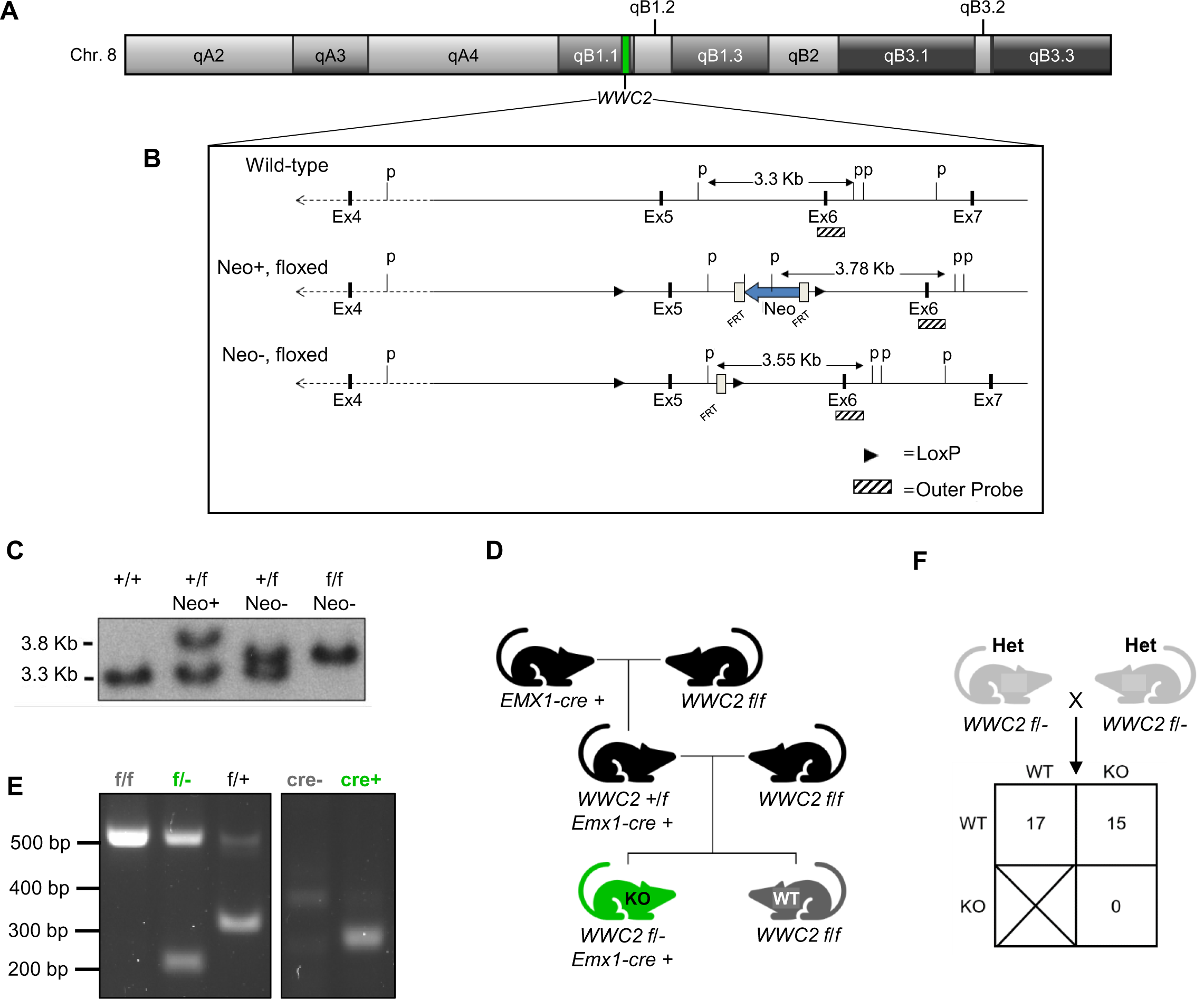
Generation of conditional *Wwc2* knockout mouse. A) Map of mouse chromosome 8, with the *Wwc2* locus highlighted in green. B) Schematic of LoxP and southern blot probe sites. C) Southern blot confirming placement of LoxP sites and resection of neomycin cassette. D) Pedigree describing generation of the WWC2 cKO and WT animals used in this study. E) Example genotyping of *Wwc2 f/f, f/-, f/+,* and *Emx1-cre* negative and positive animals. F) Mating constitutive heterozygous *Wwc2* mice never produces *Wwc2* KO pups, indicative of embryonic lethality of full body WWC2 knockout. (Chi-square test, p = 0.0008)

**Figure S3.**
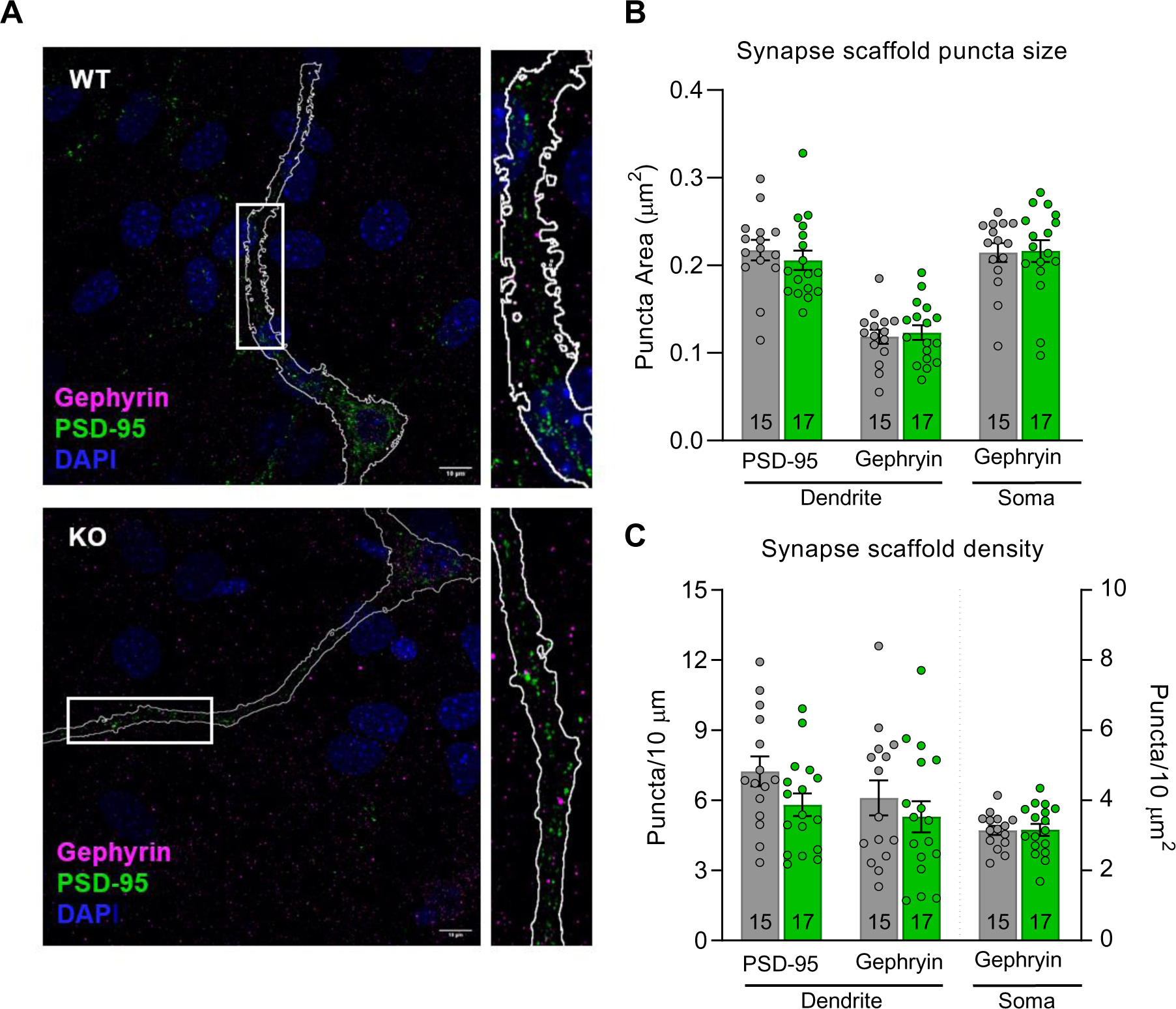
WWC2 KO does not affect distribution or size of core excitatory and inhibitory synapse scaffolds in cultured hippocampal neurons. A)Representative images of WT (top) and WWC2 cKO (bottom) neurons stained for gephryin (magenta), PSD-95 (green), and DAPI (blue). Regions of interest magnified to illustrate synapse distribution along proximal primary dendrites. Scale bars 10 µm. B,C) Unchanged excitatory (PSD-95) and inhibitory (gephryin) synapse scaffold puncta size (B) and density (C) in WWC2 KO neurons. (Puncta size; WT: PSD-95 = 0.217 ± 0.012, Gephryin_dendrite = 0.119 ± 0.008, gephryin_soma = 0.215 ± 0.011, KO: PSD-95 = 0.206 ± 0.011, Gephryin_dendrite = 0.123 ± 0.008, gephryin_soma = 0.216 ± 0.013. Puncta density; WT: PSD-95 = 7.25 ± 0.64, Gephryin_dendrite = 6.11 ± 0.747, gephryin_soma = 3.14 ± 0.13, KO: PSD-95 =5.81 ± 0.49, Gephryin_dendrite = 5.30 ± 0.67, gephryin_soma = 3.16 ± 0.17.) Biological replicates: cultures from 2 independent litters of pooled animals, n on bar graph denotes number of cells. Statistics: WT vs. KO puncta size: unpaired t-test, PSD-95 p = 0.4816, gephryin_dendrite p = 0.6289, Mann-Whitney test gephryin_soma p = 0.7944, WT vs. KO puncta density: unpaired t-test, PSD-95 p = 0.0788, gephryin_dendrite p = 0.4234, gephryin_soma p = 0.9326. Data reported as mean ± SEM.

**Figure S4.**
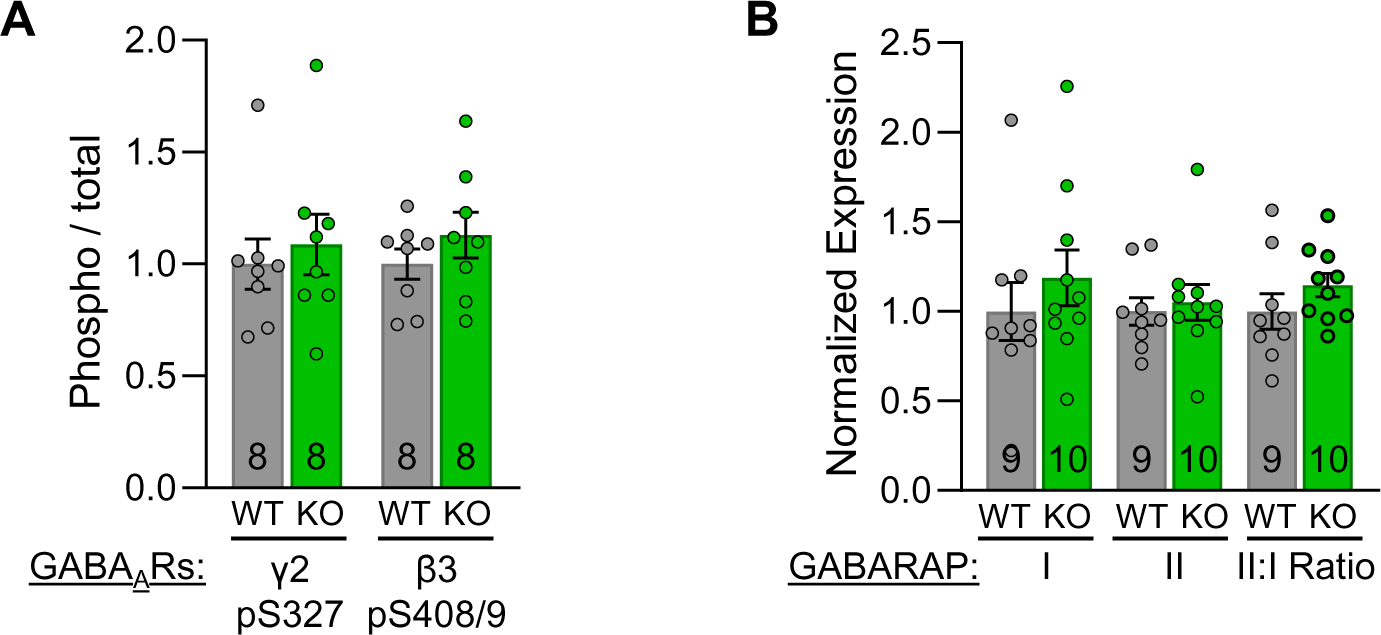
GABA_A_R γ2-S327 or β3-S408/409 phosphorylation and GABARAP activation, regulators of GABA_A_R synaptic clustering, internalization, and plasma membrane insertion, are unaffected in the WWC2 cKO hippocampus. A) Quantification of phospho to total γ2-S327 and β3-S408/409 expression assessed via western blot from hippocampal membrane-associated fraction (γ2 pS327, WT = 1.00 ± 0.11, cKO = 1.09 ± 0.14, Mann-Whitney test p = 0.07209; β3 pS408/9, WT = 1.00 ± 0.07, KO = 1.13 ± 0.10, unpaired t-test p = 0.3094). B) Loss of WWC2 did not affect GABARAP I or II expression or II/I ratio, assessed via western blot from hippocampal tissue homogenate. (GABARAP I, WT = 1.00 ± 0.16, KO = 1.19 ± 0.16, Mann-Whitney test p = 0.2775; GABARAP II, WT = 1.00 ± 0.08, KO = 1.05 ± 0.10, Mann-Whitney test p = 0.4002; GABARAP II/I ratio, WT = 1.00 ± 0.10, KO = 1.15 ± 0.07, unpaired t-test p = 0.2274). Data reported as mean ± SEM, normalized to WT (details in Methods). N indicates number of animals (A: WT = 3 males + 5 females, KO = 4 males + 4 females; B: WT = 4 males + 5 females, KO = 4 males + 6 females).

**Figure S5.**
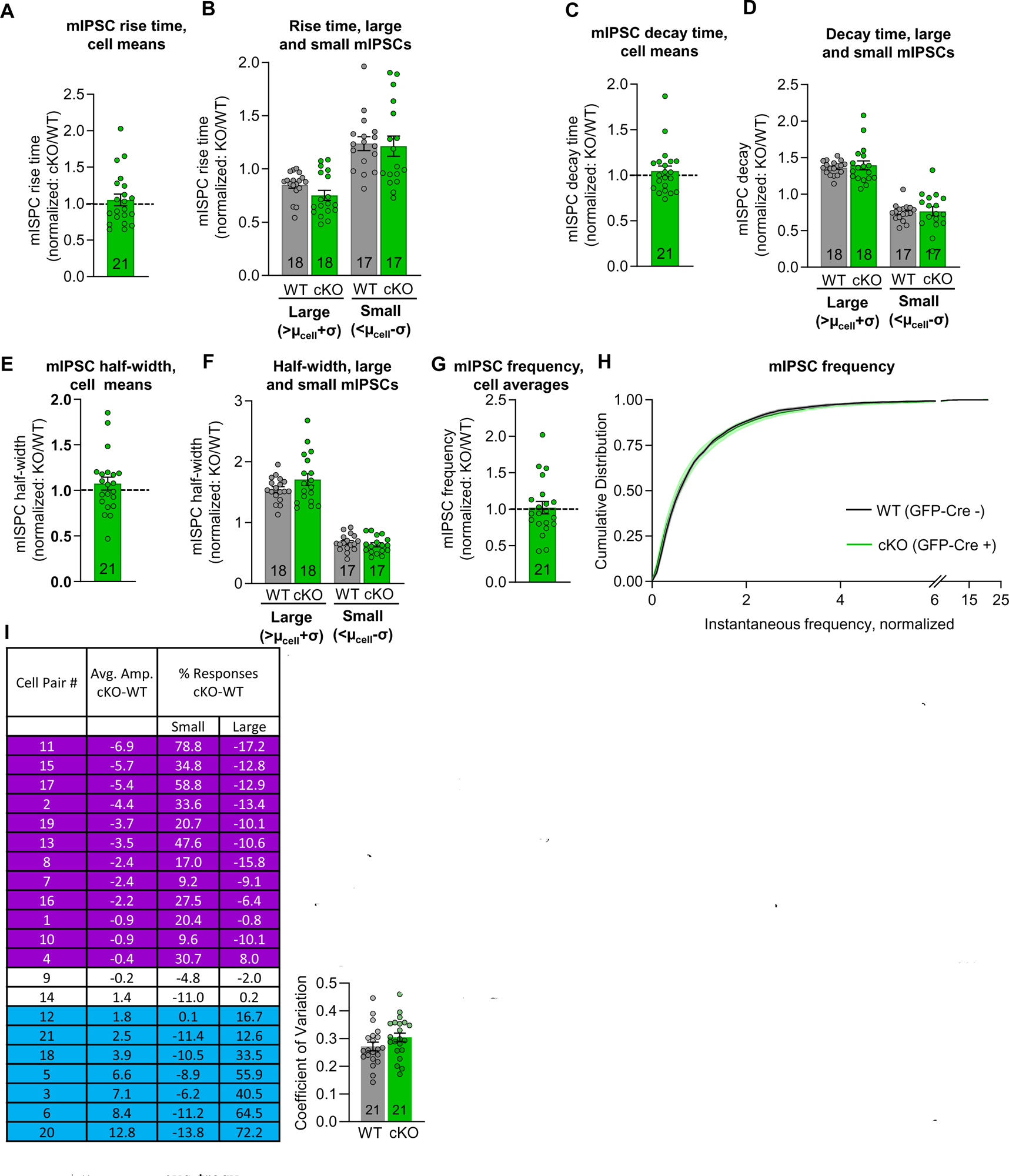
mIPSC kinetics and frequency are not altered by loss of WWC2. Cell means for mIPSC rise time (A, 10-90% rise time, cKO normalized to paired WT recording: 1.05 ± 0.08, 1-sample Wilcoxon p = 0.9457), decay time (C, 10-90% decay time, cKO normalized to paired WT recording: 1.05 ± 0.06, 1-sample Wilcoxon p = 0.8917), and half-width (E, cKO normalized to paired WT recording: 1.07 ± 0.07, 1-sample t-test p= 0.3138) are unaffected in WWC2 cKO neurons. Kinetics are also unchanged for large and small mIPSC subpopulations (B, 10-90% rise time, normalized to WT cell average, large: WT = 0.85 ± 0.03, cKO = 0.75 ± 0.05, WT vs. cKO Wilcoxon p = 0.1286, Holm-Šídák correction for multiple comparisons; small: WT = 1.24 ±0.06, cKO = 1.21 ± 0.10, WT vs. cKO Wilcoxon p = 0.7819, Holm-Šídák correction for multiple comparisons. D, 10-90% decay time, normalized to WT cell average, large: WT = 1.36 ± 0.02, cKO = 1.40 ± 0.06, WT vs. cKO Wilcoxon p = 0.9661, Holm-Šídák correction for multiple comparisons; small: WT = 0.74 ±0.03, cKO = 0.77 ± 0.06, WT vs. cKO Wilcoxon p = 0.8961, Holm-Šídák correction for multiple comparisons. F, mISPC half-width, normalized to WT cell average, large: WT = 1.55 ± 0.05, KO = 1.71 ± 0.10, WT vs. cKO paired t-test p = 0.2413 Holm-Šídák correction for multiple comparisons; small: WT = 0.67 ±0.03, cKO = 064 ± 0.03, WT vs. cKO paired t-test = 0.4702, Holm-Šídák _WT cKO_ correction for multiple comparisons.) G) Cell means for mIPSC frequency, cKO normalized to paired WT recording (1.02 ± 0.08, 1-sample t-test p = 0.8015). H) Cumulative distribution of mIPSC instantaneous frequency, normalized to paired WT recording, bin size 0.1. Data reported as mean ± SEM. N = number of cells. I) Quantification of changes in WWC2 cKO mIPSC amplitude distribution (raw cell averages). Column 1: Cell pair identity. Column 2: Change in average mIPSC amplitude in WWC2 cKO neuron (cKO-WT). Column 3: Change in percent of mIPSCs with amplitudes outside of 1 SD from the WT mean. J) Variation in mIPSC amplitude is similar in between WWC2 cKO and WT cells.s is due to cell-specific unimodal shifts in amplitude distribution. Left: Coefficient of variation calculated for raw mIPSC amplitudes is unchanged in WWC2 cKO neurons. WT (GFP-infected) = 0.27 ± 0.016, cKO (cre-infected) = 0.30 ± 0.016; p=0.14, Welch’s t-test.

**Figure S6.**
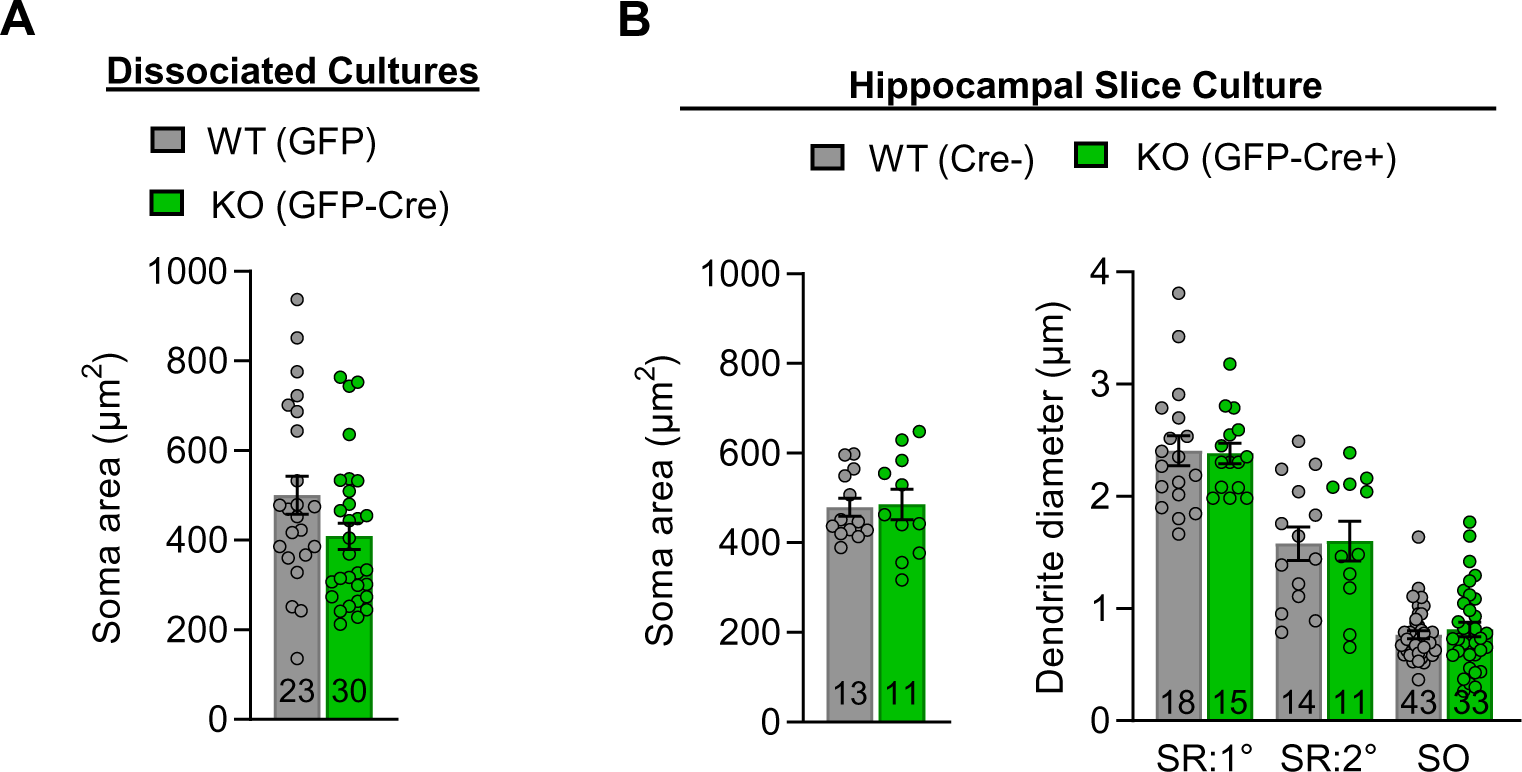
Loss of WWC2 does not affect soma size or dendritic caliber in hippocampal pyramidal cells. A) Quantification of soma area in dissociated hippocampal cultures prepared from WWC2^fl/fl^ mice. WT (GFP-infected) 500 ± 42.4, cKO (GFP-Cre-infected) 409 ± 29.4, Mann Whitney test p = 0.0858. N indicates number of cells from 2 independent cultures. Center: Quantification of soma surface area in organotypic hippocampal slice cultures prepared from WWC2^fl/fl^ mice. WT (untransfected) 479 ± 20.4, cKO (GFP-Cre-transfected) 486 ± 33.7, unpaired t-test p = 0.8681. B) Quantification of stratum radiatum (SR, primary and secondary) and stratum oriens (SO) dendritic diameter in CA1 pyramidal cells from organotypic slice cultures used for mPSC recordings (1° SR: WT = 2.41 ± 0.13, cKO = 2.38 ± 0.09, Mann-Whitney with Holm-Šídák correction for multiple comparisons, p = 0.9856. 2° SR: WT = 1.58 ± 0.15, cKO = 1.60 ± 0.18, Mann-Whitney with Holm-Šídák correction for multiple comparisons, p = 0.9856. SO: WT = 0.77 ± 0.03, cKO = 0.81 ± 0.07, Mann-Whitney with Holm-Šídák correction for multiple comparisons, p = 0.9856.) Data reported as mean ± SEM. N indicates number of cells for soma area and number of dendritic segments for dendritic caliber (18 WT and 15 KO cells analyzed). Data from 3 separate litters.

## Key resources table

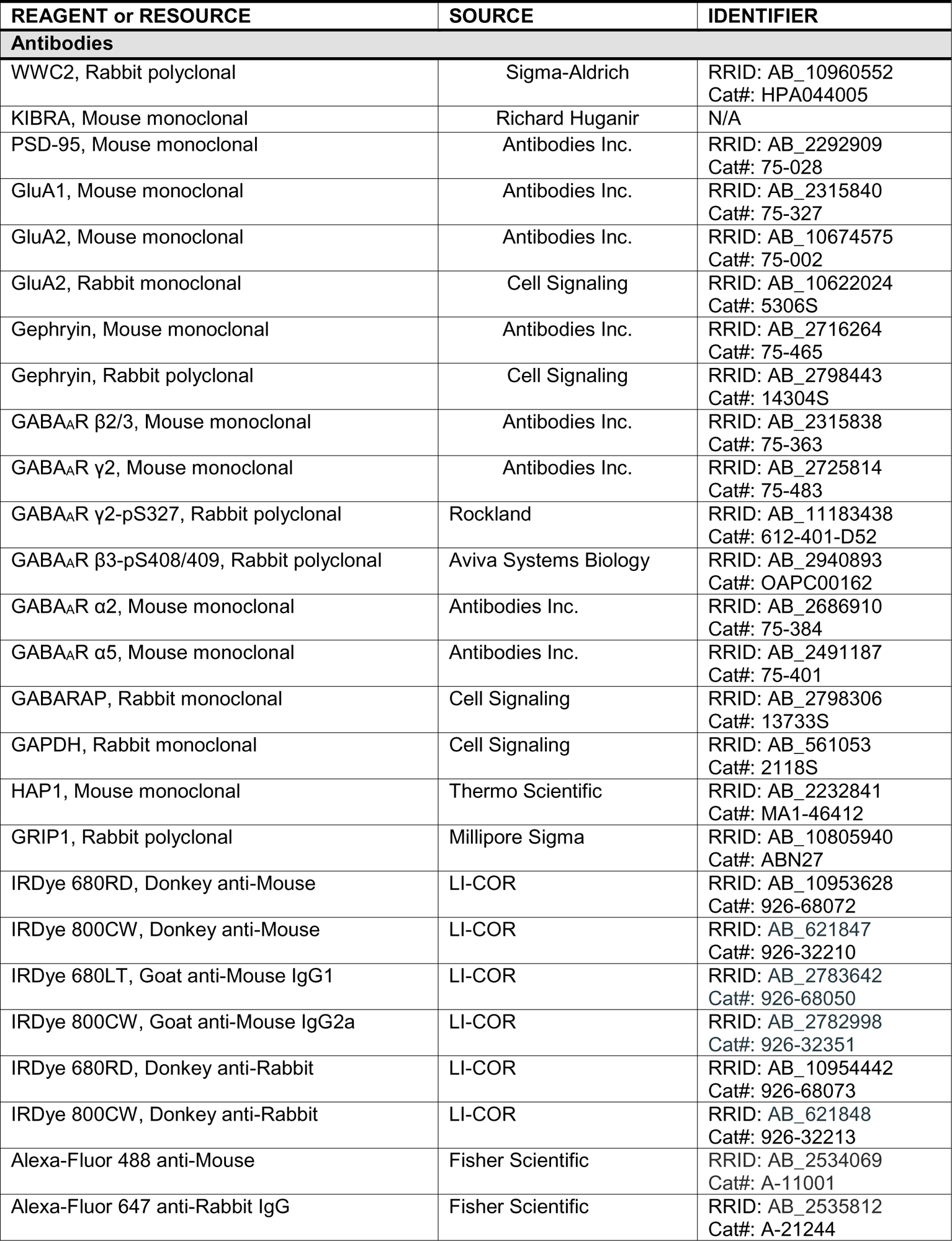
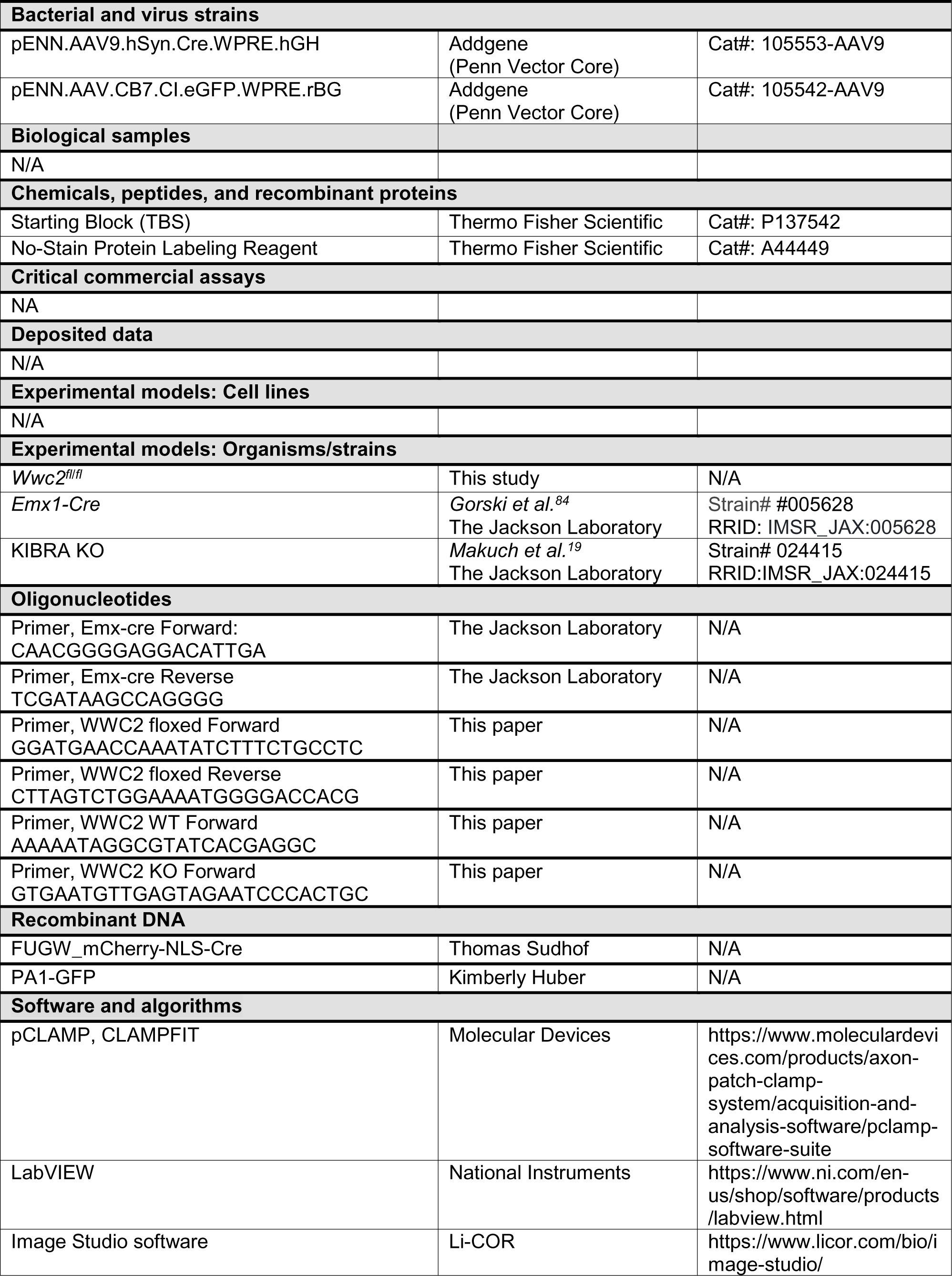
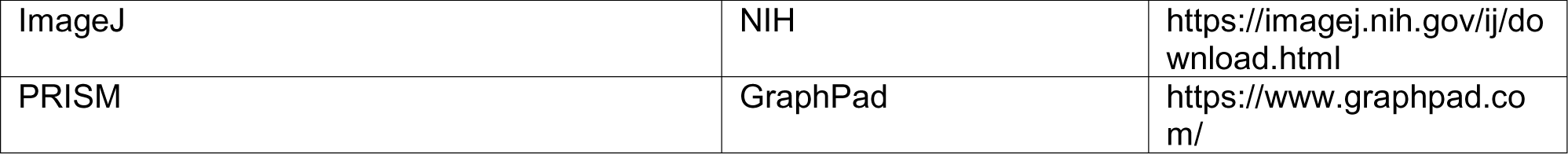

